# Fermentation-Derived Metabolites Shape Host Biology to Attenuate Severity of Inflammatory and Metabolic Disease

**DOI:** 10.64898/2026.07.30.741534

**Authors:** Elisa B. Caffrey, Elektra Kantzari Robinson, Jessica L. Fessler, Anita Reddy, Samantha G. Hernandez, Yifan Gao, Emma R. Guiberson, Sean P. Spencer, Erica D. Sonnenburg, Justin L. Sonnenburg

## Abstract

Fermented foods are among the few dietary interventions shown to increase gut microbiome diversity and reduce systemic inflammation in healthy adults, yet the underlying mechanisms remain poorly defined. Metabolites produced during food fermentation, termed fermentation-derived metabolites (FDMs), represent a largely uncharacterized pool of bioactive compounds that may directly mediate the physiological effects of fermented food consumption. Here, we characterize the metabolite landscape of ten vegetable-based fermented foods using metabolomics, identifying conserved enrichment of aromatic and branched-chain amino acid derivatives across diverse substrates. Using sauerkraut as a chemically representative model system, we show that metabolite extracts from wild green sauerkraut (wGS-FDMs) remodel intestinal and systemic immune populations and shift gut microbiome composition in healthy mice. wGS-FDMs suppressed NF-κB activation and pro-inflammatory cytokine secretion in vitro and decreased colitis severity in vivo. In a chronic high-fat diet model, wGS-FDMs attenuated weight gain and improved glucose and insulin tolerance, consistent with stimulation of GLP-1 secretion in vitro. Collectively, these findings establish FDMs as biologically potent dietary components capable of simultaneously modulating immune, microbial, and metabolic homeostasis across multiple physiological systems, positioning metabolites from fermented foods as an important and underappreciated class of dietary effectors in the context of chronic disease.

## Introduction

The gut microbiome (or gut microbiota) is recognized as a central key in host physiology, influencing immune tone, metabolic homeostasis, and susceptibility to chronic disease^1–3^. A critical mediator of microbiota-host crosstalk is microbial metabolism, where anaerobic fermentation of dietary substrates generate a diverse pool of microbiota-derived metabolites (MDMs) that can act on host epithelial cells^4,5^, immune cells^6^, and members of the gut microbiota^7,8^, and can be absorbed to enter systemic circulation, mediating microbial-host interactions^9^. Many MDMs, including short-chain fatty acids (SCFAs) and aromatic amino acid metabolites, exert profound effects across epithelial, immune, and neuronal compartments^7,10^. As research on the gut microbiome has grown, metabolites have emerged as key for developing mechanistic insights into the gut microbiota and their role in both health and disease^11–15^.

It is widely understood that diet is a dominant driver of microbiome composition and function^16–18^, and that gut microbiota diversity of industrialized populations is lower relative to non-industrialized cohorts^19–24^ with higher diversity correlated with improved metabolic parameters and immune profiles^22^. Probiotic consumption, fecal microbiota transplantation (FMT), and dietary modifications have been used to alter gut microbiome composition, yet a dietary intervention that increases microbiome diversity in healthy adults living in an industrialized lifestyle has remained elusive^25–28^. To date, fermented food-rich diets are the only dietary intervention shown to increase gut microbiome diversity and decrease circulating markers of inflammation in industrialized populations^29^, positioning fermented foods as potent and clinically meaningful modifiers of host immune state. Beyond their effects on microbial ecology, fermented foods deliver a rich milieu of metabolites produced by microbes during fermentation. The rapid expansion of understanding how MDMs impact host biology in numerous ways positions the metabolites generated during food fermentation as compelling and underexplored host modulators that may operate independently of live microorganisms and complement the exogenous pools of MDMs that regulate host physiology.

Fermentation is a metabolic process central to both gut microbiome ecology and food production. Fermented foods introduce an additional, exogenous pool of structurally analogous metabolites, referred to as fermentation-derived metabolites (FDMs), that may mimic or complement endogenous microbial metabolism and impact host biology^12^. Recent studies have begun to characterize FDMs produced during food fermentation^30–34^, but the breadth, variability, and functional consequences remain to be understood. A growing body of work suggests that metabolites, rather than the live microbes themselves, may underlie many of the physiological effects attributed to fermented foods. Several recent studies have demonstrated that fermented foods can confer measurable host effects when microbial viability is removed. For example, Nielsen et al. showed that sauerkraut improved IBS symptoms independent of pasteurization^35^; while Schropp et al. reported that consumption of pasteurized sauerkraut increased circulating butyrate in participants when compared to participants consuming unpasteurized sauerkraut, with both groups showing improved inflammatory tone^36^. Together, these findings point toward FDMs as both tractable and biologically significant, motivating a metabolite-centered approach to understanding fermented food function.

We followed a reverse translation framework, building upon the human clinical observations reported by Wastyk et al.^29^. We focused specifically on vegetable-based fermented foods (VBFFs), which are associated with increased microbial ASVs, and whose simpler food matrix relative to dairy enables FDMs to be systematically characterized. VBFFs provide experimentally tractable models for dissecting the mechanisms of microbial community assembly ^37,38^, and clinical intervention studies have demonstrated that their consumption increases gut microbiome diversity in a dose-dependent manner^29^. Using semi-targeted and targeted metabolomics, we defined the chemical landscape of VBFF and found wild green sauerkraut to be a chemically representative and experimentally tractable model system for studying FDMs. Exposure to wild green sauerkraut metabolite extracts (wGS-FDM) remodeled the intestinal, microbial, and systemic immune landscape under homeostatic conditions and conferred protection in both an acute inflammatory model of DSS-induced colitis and a chronic inflammatory model of high-fat diet. wGS-FDMs suppressed macrophage NF-κB activation and TNF-α secretion and stimulated enteroendocrine GLP-1 secretion in vitro, demonstrating direct bioactivity across immune and endocrine cell types. Together, these findings position FDMs as biologically potent dietary bioactives with broad implications in the prevention and management of metabolic and inflammation-driven disease.

## Results

### Vegetable ferments share a conserved group of metabolites

Mapping the metabolite landscape of FDMs remains a challenge for the mechanistic exploration of fermented foods, microbial community dynamics, and potential microbial-host interactions. To understand the role of microbial activity on the fermented food metabolome, we employed a previously validated semi-targeted method for identification of gut microbiome-derived metabolites^39,40^ and a short chain fatty acid (SCFA)-targeted method using liquid chromatography-mass spectrometry (LC-MS) longitudinally across ten cabbage-based ferments: wild baechu kimchi with fish sauce (wBK), wild vegan baechu kimchi (wVBK), baechu kimchi with kimchi starter and fish sauce (sBK), vegan baechu kimchi with starter (sVBK), nabak kimchi with (sNK) and without (wNK) starter, curtido (wC), red cabbage sauerkraut (wRS), green cabbage sauerkraut with starter (sGS), and without starter (wGS) (see methods) (Fig. 1A). Samples were collected across multiple timepoints of fermentation, with the final timepoint defined based on pH (Supplementary Table 1).

**Figure 1.**
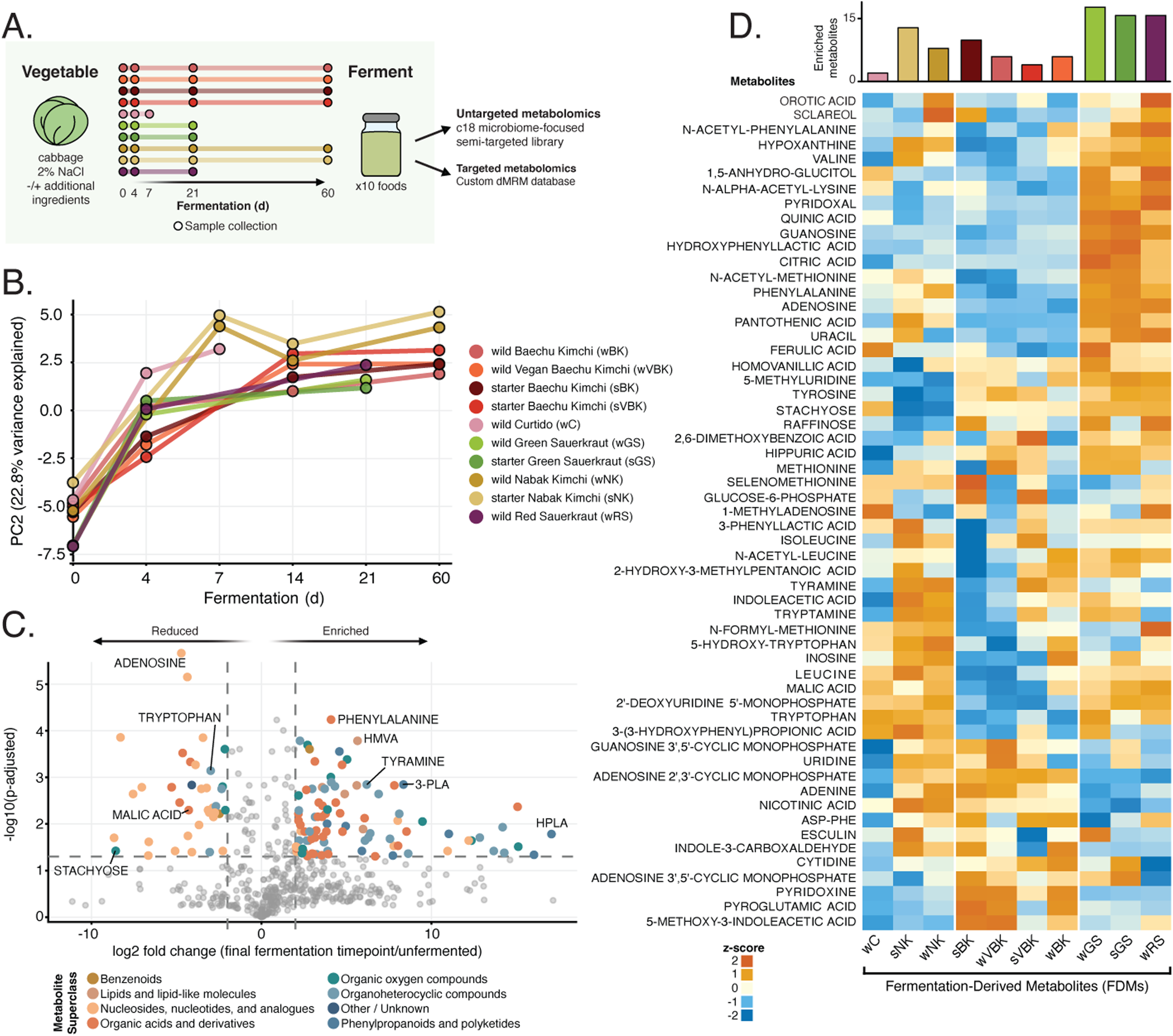
Wild fermented green sauerkraut (wGS) develops a uniquely enriched bioactive metabolite profile among ten fermented vegetables. (A) Experimental design. Ten fermented vegetable preparations were sampled across a fermentation time course (days 0, 4, 7, 21, 60), using a bacterial starter (s) or wild starter (w), and analyzed by untargeted C18 semi-targeted metabolomics and targeted dMRM. (B) Probabilistic principal component analysis (PC2, 22.8% variance explained vs fermentation time) of metabolite profiles across all ten foods (w = wild fermentation) (s = starter fermentation used) and (V = vegan). (C) Volcano plot of within-food metabolite changes (fermented vs. day 0, BH-adjusted p-value); all foods pooled. Enriched metabolites include amino acids, phenylpropanoids (3-phenyllactic acid (3-PLA), 4-hydroxyphenyllactic acid (HPLA)) and branched-chain hydroxy acids (2-hydroxy-3-methylpentanoic acid (HMVA)); depleted metabolites include adenosine, stachyose, and malic acid. Points colored by metabolite superclass. Dashed lines indicate significance and fold-change thresholds. (D) Heatmap of metabolite abundance across all foods and timepoints. Wild baechu kimchi with fish sauce (wBK), wild vegan baechu kimchi (wVBK), baechu kimchi with kimchi starter and fish sauce (sBK), vegan baechu kimchi with starter (sVBK), nabak kimchi with (sNK) and without (wNK) starter, curtido (wC), red cabbage sauerkraut (wRS), green cabbage sauerkraut with starter (sGS), and without starter (wGS). The top bar chart shows frequency of high-abundance metabolites (scale > 1) per food.

Untargeted metabolomics revealed chemical diversity across the ferments (643 features, mean 483 per sample, range of detected features 313-622) (Supplementary Table 2), with baechu kimchis being distinct from vegetable ferments made with fewer ingredients (Supplementary Fig. 1A). Metabolite composition shifts progressively with fermentation time with broad similarities evident in a principal component analysis, a progression driven by production and depletion of metabolites associated with fermentation (Fig. 1B, Supplementary Fig. 1B)^7,9,31,41–44^. Across all fermented foods tested, nucleosides, nucleotides, and related analogues were broadly depleted, consistent with microbial uptake during active growth^12,45^. In contrast, organoheterocyclic and phenylpropanoid compounds were consistently enriched, revealing conserved chemical trajectories of lactic acid fermentation (Fig. 1C). Notably, we observed robust increases in branched-chain amino acid derivatives, including 2-hydroxy-3-methylpentanoic acid (HMVA), as well as aromatic amino acid metabolites, such as 3-phenyllactic acid (3-PLA) and 4-hydroxyphenyllactic acid (4-HPLA), highlighting shared microbial transformations across substrates (Supplementary Fig. 2A-D). Short-chain fatty acid (SCFA) quantification confirmed the expected production of acetate and lactate across conditions, consistent with active lactic acid fermentation (Supplementary Fig. 2E). Notably, inspection of unassigned features revealed increased production of N-lactoyl-phenylalanine (Lac-Phe), recently associated with beneficial metabolic effects of exercise^46^, underscoring the broader discovery potential of fermented food chemical space (Supplementary Fig. 3A-B). Together these findings motivate a metabolite-centered framework for understanding fermented food impact on humans.

Analysis of the annotated semi-targeted metabolites further highlighted substrate-specific profiles (Fig. 1D). Notably, wild fermented green sauerkraut (wGS) exhibited a chemically rich metabolite profile with a metabolome most representative of other vegetable ferments (Supplementary Fig. 4A-B). Thus, we select wGS as a model to probe the impact of vegetable-fermentation-derived metabolites on health.

### wGS-FDMs reshape the gut physiology, gut microbiome, and immune landscape in vivo

Given that fermented food consumption decreases circulating markers of inflammation in healthy adults^29^, we investigated whether metabolites extracted from wild green sauerkraut fermented for 3 weeks (wGS-FDMs) could modulate gut physiology and immune homeostasis. C57BL/6 mice harboring a conventional microbiome were treated with wGS-FDMs reconstituted in their drinking water. After 17 days intestinal morphology, microbiome composition, and immune parameters were assessed (Fig. 2A). wGS-FDM treatment increased large intestinal length (Fig. 2B), prompting us to examine whether epithelial proliferation underlies this effect. Flow cytometry was used to quantify rapidly dividing Ki67^+^ epithelial cells (EpCAM^+^). Ki67^+^EpCAM^+^ cell frequency was unchanged in the large intestine of wGS-FDMs-treated mice (Supplementary Fig. 5A). However, small intestinal epithelial proliferation was elevated (Supplementary Fig. 5B), without changes in MHCII expression indicating that increased proliferation occurred in the absence of epithelial inflammation (Supplementary Fig. 5C, 6). Histological analysis supported epithelial proliferation with greater small intestinal villus height in wGS-FDM-treated animals (Supplementary Fig. 5D-F), demonstrating that short-term exposure of fermented food metabolites is sufficient to remodel the intestinal epithelium.

**Figure 2.**
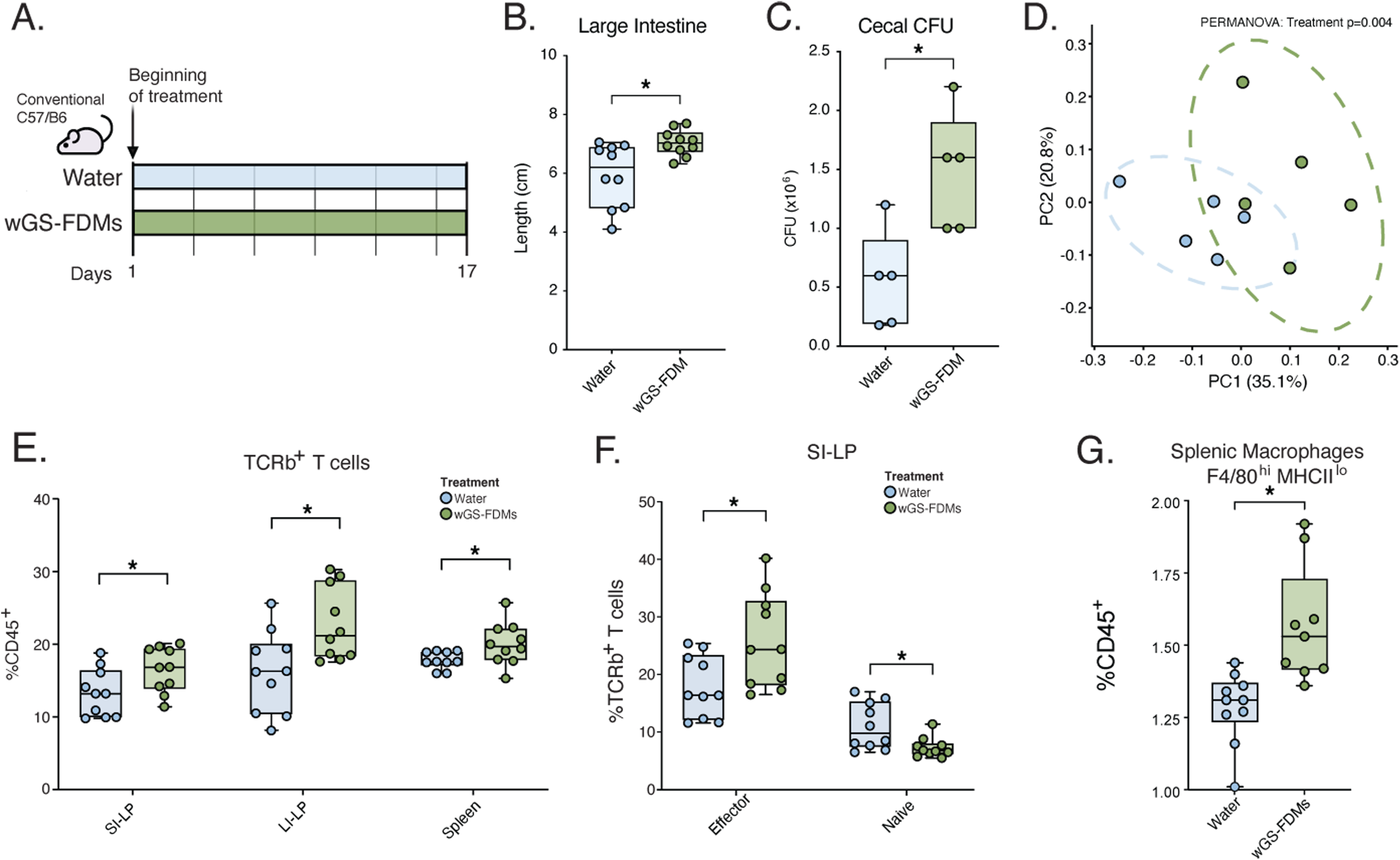
wGS-FDMs reshape intestinal physiology, microbiome composition, and immune populations in conventional mice. (A) Experimental design. C57BL/6 SPF mice received water or wGS-FDMs for 17 days. Gut morphology and microbiome load: large intestinal length (B), cecal colony forming units (CFU) (C). (D) Principal coordinates analysis (PCoA) of Bray-Curtis dissimilarities at the species level, with water (blue) and wGS-FDM (green) treatment groups across PC1 (33.7%) and PC2 (20.1%). (E) Frequency of TCRb+ T cells among CD45+ live cells in small intestinal lamina propria (SI-LP), large intestinal lamina propria (LI-LP), and spleen. (F) Frequency of effector or naive TCRb+ T cells in the small intestinal lamina propria. (G) Frequency of splenic F4/80hiMHCIIlo (resident) macrophages. (H) Schematic of the NFκB reporter macrophage assay. (I) Cell viability at 200ng/ml lipopolysaccharide (LPS) expressed as percentage of untreated control (CTL) at 0 ng/mL LPS; conditions tested: CTL, lithocholic acid (LCA), cabbage metabolite (CB-M), 24hr-wGS-FDM, 96hr-wGS-FDM, 1w-wGS-FDM, and 3w-wGS-FDM. (J) NFκB-GFP reporter fluorescence (GFP mean fluorescence intensity (MFI)) normalized to CTL at 200 ng/mL LPS across all conditions. Statistical comparisons by Mann-Whitney U or unpaired t-test as appropriate. Data represent mean ± SEM; n = [10] mice per group. *p<0.05, **p<0.01, ***p<0.001.

Given that intestinal remodeling can reflect changes in the microbial environment, we examined whether wGS-FDMs alter bacterial load and microbiome composition. Cecal CFUs were elevated in wGS-FDMs-treated mice (Fig. 2C), indicating short-term fermented food metabolite exposure increases luminal microbial load. wGS-FDM-treated mice exhibit a distinct cecal microbiome composition relative to controls, evidenced by metagenomic sequencing (Fig. 2D). Notably, wGS-FDM treatment enriched the relative abundance of *Akkermansia muciniphila*, a mucus-associated commensal, (Supplementary Fig. 7A-C) showing that fermentation-derived metabolites (i.e., without the live bacteria or food substrate) are sufficient to reshape microbiota composition.

We next examined whether wGS-FDM treatment reshaped immune populations using flow cytometry of cells from intestinal and peripheral compartments. In the small intestinal lamina propria (SI-LP), large intestinal lamina propria (LI-LP) and spleen, wGS-FDM-treated mice had increased TCRβ^+^ T cells as a percentage of total immune cells (CD45^+^)(Fig. 2E). Phenotypic analysis of the expanded SI-LP T cell pool revealed a shift in composition, with a reduction in naive T cells and a corresponding increase in effector T cells, indicating that wGS-FDMs increase the proportion of antigen-experienced cells (Fig. 2F). This T cell compositional shift was not observed in the LI-LP or spleen (Supplementary Fig. 8A-B), consistent with the distinct immunological environments and antigen exposure profiles of these compartments. In the spleen, wGS-FDM treatment was associated with an increase in the proportion of tissue-resident macrophages (F4/80^hi^MHCII^lo^) relative to untreated controls (Fig. 2G). This population displayed elevated CD80 surface expression in wGS-FDM-treated mice (Supplementary Fig. 8C-D), suggesting increased co-stimulatory potential within an otherwise tolerogenic macrophage subset.

To determine whether the immunological effects of wGS-FDMs require an intact microbiota, we administered wGS-FDMs to germ-free C57BL/6 mice for 17 days (Supplemental Fig. 9A). Despite recapitulating intestinal epithelial proliferation seen in conventional mice (Supplementary Fig. 9B-C), wGS-FDM failed to alter TCRβ^+^ T cell proportions in the SI-LP, LI-LP, or spleen (Supplementary Fig. 9D-F), or to affect splenic macrophage activation state or CD80 expression (Supplemental Fig. 9G-H). These findings dissociate direct epithelial effects of wGS-FDMs from their immune modulatory activity and implicate microbiota-dependent signals in wGS-FDM-driven adaptive and innate immune remodeling. Together, these findings demonstrate that wGS-FDMs orchestrate coordinated, microbiota-dependent remodeling of the intestinal and systemic immune landscape under homeostatic conditions, without inducing overt inflammation.

### wGS-FDMs protect against DSS-induced colitis and associated microbiome disruption

Given that wGS-FDMs remodel immune tone and epithelial architecture under homeostatic conditions, we next sought to determine whether these metabolites could confer protection in the context of overt intestinal inflammation using the DSS-induced colitis model. Mice were treated with the 3-week wGS-FDM for seven days prior to DSS exposure and continued daily dosing throughout the experiment (Fig. 3A). wGS-FDM-treated mice showed attenuated weight loss during DSS challenge and improved recovery during the resolution phase relative to water controls (Fig. 3B). Disease activity index (DAI) scores, a composite measure of weight loss, stool consistency, and rectal bleeding, were also lower in wGS-FDM-treated mice across the DSS and early recovery periods (Fig. 3C). At the experimental endpoint, wGS-FDM-treated mice had increased colon length, greater cecal mass, and higher colonic *Cldn3* expression, consistent with reduced epithelial damage and improved intestinal integrity (Figs. 3D-E; Supplementary Fig. 10A). Stool 16S rRNA sequencing revealed that wGS-FDM-treated mice harbored a compositionally distinct microbiome relative to water controls (Supplementary Fig. 10B) and were protected from DSS-associated microbiome disruption, as reflected by reduced within-mouse Bray-Curtis dissimilarity between Week 0 and Week 3 (Fig. 3F).

**Figure 3.**
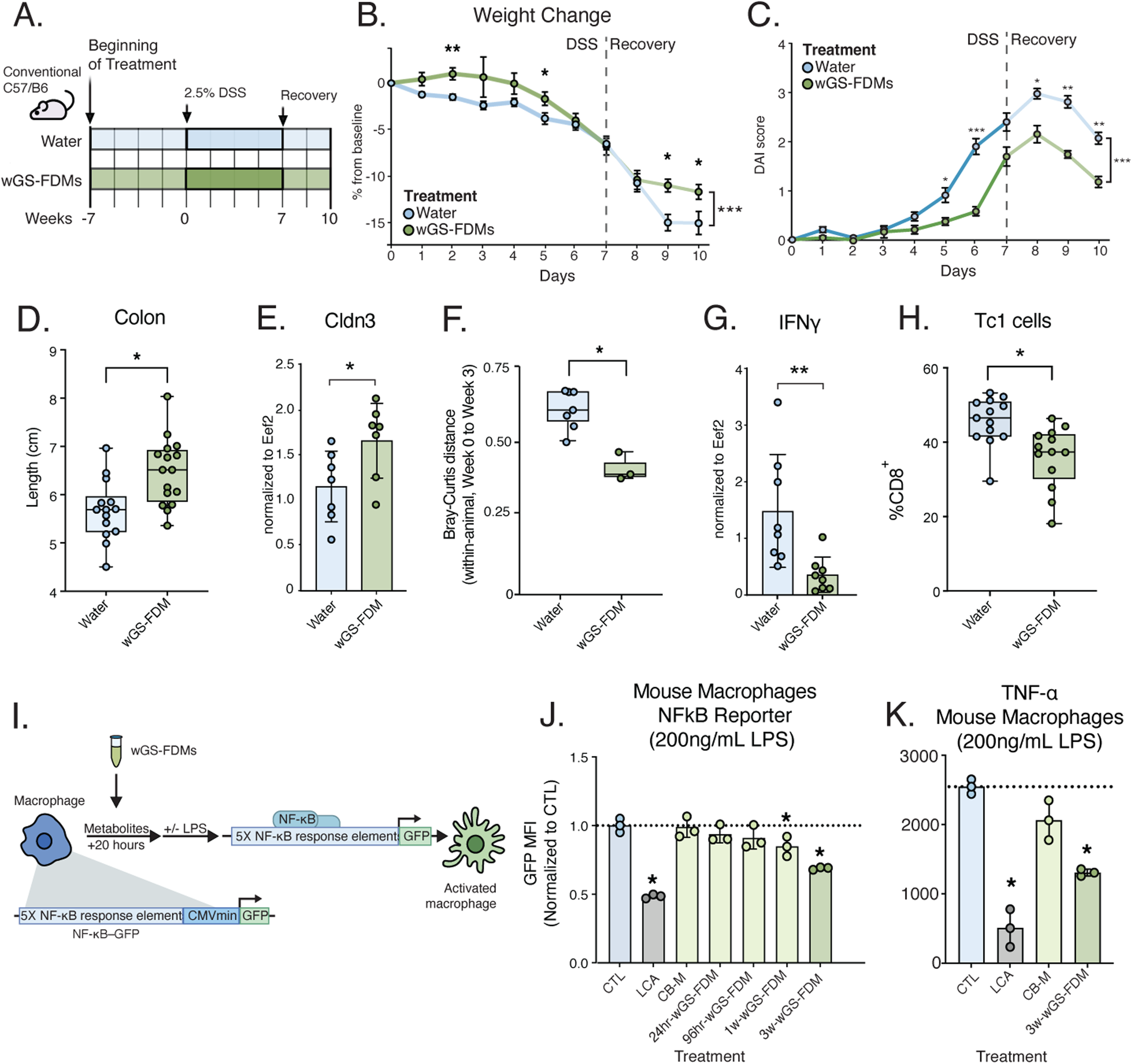
wGS-FDMs attenuate DSS-induced colitis and suppress LPS-driven NF-κB activation in macrophages. (A) Schematic of the dextran sodium sulfate (DSS) induced colitis experimental design. (B) Body weight expressed as percentage change from baseline over the course of the experiment; the DSS and recovery phases are indicated (two-way ANOVA, *p-value* < 0.05) (C) Disease Activity Index (DAI) score over time. (D) Large intestinal length (cm) at experimental endpoint (day 10). (E) RT-qPCR of Claudin-3 (Cldn3) from large intestinal tissue normalized to Eukaryotic Translation Elongation Factor 2 (Eef2). (F) Bray-Curtis dissimilarity between paired stool samples (Week 0 to Week 3) within each mouse (Wilcoxon rank-sum, *p-val = 0.043*) (G) RT-qPCR of Interferon-gamma (IFN-γ) from large intestinal tissue normalized to Eef2. (H) Frequency of Tc1 cells among CD8+ cells in the large intestinal lamina propria. Data are shown as mean ± SEM. Statistical comparisons by Mann-Whitney U test or unpaired t-test. *In vitro* experiments performed in 2 independent replicates. n=12 mice per group in vivo. *p<0.05, **p<0.01, ***p<0.001. ns, not significant.

Immune profiling of large intestinal tissue revealed lower *Ifng* expression in wGS-FDM-treated colitic mice as measured by RT-qPCR, indicating attenuated pro-inflammatory cytokine signaling (Figs. 3G). Flow cytometric analysis of colonic CD8^+^ T cells showed a smaller proportion of the IFN-γ producing Tbet^+^ population (Tc1) in wGS-FDM-treated mice at three weeks (Fig. 3H), consistent with dampened cytotoxic T cell-driven tissue inflammation. Together, these data indicate that wGS-FDMs mitigate inflammatory barrier damage, preserve microbiome stability, and promote epithelial repair during acute colitis, consistent with the *in vitro* suppression of innate immune signaling.

To interrogate the cellular mechanism underlying the observed *in vivo* protection, we next examined whether wGS-FDMs could directly suppress innate immune signaling using an NFκB reporter macrophage model. Metabolites extracted across the course of wGS fermentation (0 hours, 24 hours, 96 hours, 1 week, and 3 weeks) were applied to LPS-stimulated immortalized murine macrophages (Fig. 3I). wGS-FDMs suppressed NF-κB reporter activity at later fermentation stages (1 and 3 weeks) (Fig. 3J). Importantly, no reduction in cell viability was observed, and NF-κB reporter activity was unchanged in the absence of LPS stimulation, confirming that wGS-FDM effects are not driven by cytotoxicity or constitutive immune suppression (Supplementary Fig. 11A-B). Correspondingly, 3-week wGS-FDMs reduced Tnfα (Fig. 3K) and Il6 (Supplementary Fig. 11C) expression in macrophages compared to unfermented cabbage metabolite extract (CB-M) controls, with no changes at baseline (Supplementary Fig. 11D-E). These findings were also observed in human THP-1 cells (Supplementary Fig. 11F). Together, these findings establish that fermentation-derived metabolites are sufficient to confer anti-inflammatory bioactivity, suppressing macrophage inflammatory tone across both mouse and human systems.

### wGS-FDMs enhance GLP-1 secretion and improve immune and metabolic outcomes in a high-fat diet model

Having demonstrated that wGS-FDMs suppress acute mucosal inflammation, we next asked whether these effects extend to chronic inflammation using a high-fat diet model of obesity-associated metabolic disease. Conventional C57BL/6 mice were placed on a high-fat diet (HFD) and were either administered water (control) or wGS-FDMs treated water for 9 weeks (Fig. 4A). wGS-FDM-treated mice showed attenuation of HFD-induced weight gain beginning at week 1 and persisting through the end of the experiment (week 9) (Fig. 4B, two-way ANOVA, p < 0.001). wGS-FDM-treated mice also exhibited higher cecal weight and cecal CFU relative to untreated controls (Supplementary Figs. 12A-B), consistent with wGS-FDM-driven remodeling of the intestinal microenvironment observed under healthy conditions (Fig. 2C).

**Figure 4.**
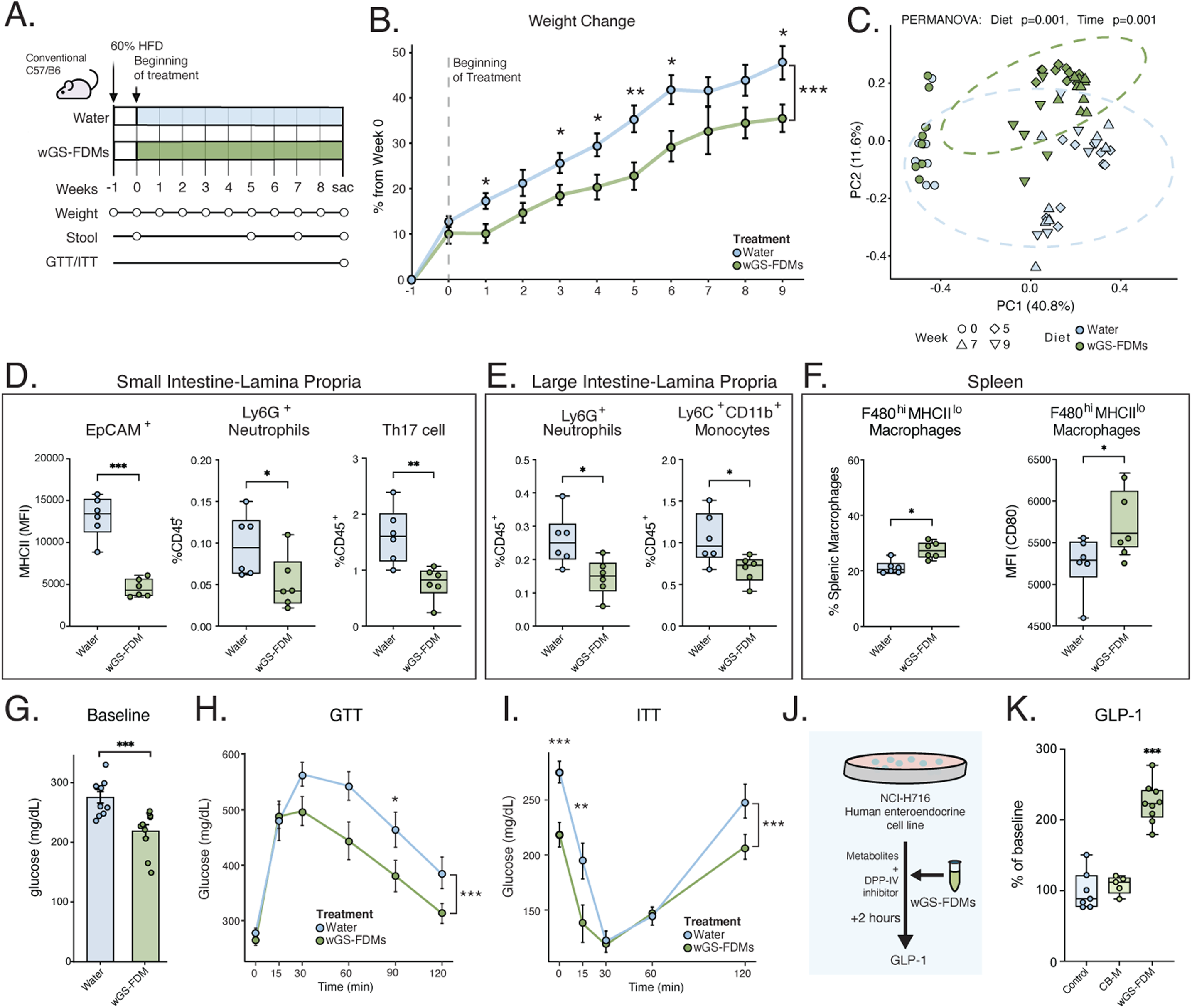
wGS-FDMs attenuate HFD-induced weight gain, inflammation, and metabolic dysfunction while stimulating GLP-1 secretion. (A) Schematic of the experimental design. C57BL/6 conventional mice were fed a 60% high-fat diet (HFD) and received daily oral gavage of water or wGS-FDMs (300µL) for 10 weeks. (B) Body weight expressed as a percentage change from week 0 over the course of 10 weeks. The dashed line marks the beginning of treatment. Per-timepoint asterisks: Welch’s t-test (*p < 0.05); Mixed repeated-measures ANOVA (Treatment: *p-value* = 0.019; Time *p-value* < 0.0001; Treatment × Time p = 0.497) (C) Principal coordinates analysis (PCoA) of Bray-Curtis dissimilarities at the ASV level in stool samples from HFD-fed water and wGS-FDM-treated mice across weeks 0, 5, 7, and 11, with dashed ellipses representing 95% confidence intervals per treatment group; PERMANOVA revealed significant effects of both diet (p = 0.001) and time (p = 0.001). (D) Flow cytometric analysis of the small intestine lamina propria from water and wGS-FDM-treated mice. Major Histocompatibility Complex Class II (MHCII) expression (mean fluorescence intensity (MFI)) on EpCAM⁺ epithelial cells, frequency of Ly6G⁺ neutrophils among CD45⁺ cells, and frequency of Th17 cells among CD45⁺ cells are shown. (E) Analysis of the large intestine lamina propria. Frequency of Ly6G⁺ neutrophils and Ly6C⁺CD11b⁺ monocytes among CD45⁺ cells are shown for water and wGS-FDM-treated mice. (F) Splenic immune cell analysis. The percentage of F480ʰⁱMHCIIˡᵒ macrophages among total splenic macrophages (left) and MHCII (CD80) MFI on F480ʰⁱMHCIIˡᵒ macrophages (right) are shown. (G) Fasting blood glucose (mg/dL) at week 9. (H) Blood glucose (mg/dL) during an oral glucose tolerance test (GTT) at 0, 15, 30, 60, 90, and 120 minutes following glucose administration (mixed ANOVA *p-val* < 0.05). (I) Blood glucose (mg/dL) during an intraperitoneal insulin tolerance test (ITT) at 0, 15, 30, 60, and 120 minutes following insulin administration (mixed ANOVA *p-val* < 0.001). (J) Schematic of the glucagon-like peptide-1 (GLP-1) secretion assay. H-716 human enteroendocrine cells were incubated with wGS-FDMs and a Dipeptidyl peptidase (DPP-IV) inhibitor for 2 hours in the presence of 100mM glucose. (K) GLP-1 secretion expressed as percentage of baseline. Conditions: 100mM glucose alone (water control; CTL), cabbage metabolites (CB-M) and wGS-FDMs. Data are shown as mean ± SEM. Statistical comparisons by Mann-Whitney U test or unpaired t-test. n = [6] mice per group; GLP-1 assay performed in 3 independent experiments. *p<0.05, **p<0.01, ***p<0.001. ns, not significant.

16S rRNA sequencing of stool revealed differences in microbiota composition between wGS-FDM- and water-treated HFD mice (Fig. 4C). Differential abundance analysis identified multiple taxa enriched in wGS-FDM-treated mice, including several Lachnospiraceae family members, taxa associated with short-chain fatty acid production and metabolic homeostasis, while Lachnoclostridium, Sporosarcina, and Jeotgalicoccus were elevated in water controls (Supplementary Figs. 12C-D), suggesting wGS-FDMs selectively promote a microbiota configuration associated with metabolic benefit^47–49^. Metagenomic comparisons of treated and untreated baseline and HFD mice showed a shift in the composition of the treated HFD group towards the microbiome of conventional diet mice, further suggesting that wGS-FDMs mitigate the impact of HFD through the microbiome (Supplementary Fig. 13A-B).

Beyond the microbiome, wGS-FDM treatment broadly reduced inflammatory immune infiltration across intestinal compartments. In the SI-LP, wGS-FDM-treated HFD mice exhibited reduced EpCAM^+^ cell MHCII expression, Ly6G^+^ neutrophils, and Th17 cells (Fig. 4D). In the LI-LP, Ly6G^+^ neutrophils and Ly6C^+^CD11b^+^ monocytes were similarly reduced (Fig. 4E). Systemically, splenic F4/80^hi^MHCII^lo^ macrophage frequency and CD80 MFI were both elevated in wGS-FDM-treated mice relative to water controls (Fig. 4F), mirroring the immunological pattern observed in healthy animals and suggesting a conserved effect of wGS-FDMs on macrophage co-stimulatory tone across inflammatory contexts.

Accompanying these immune changes, wGS-FDM-treated HFD mice displayed improved metabolic parameters, including a lower fasting blood glucose (Fig. 4G), enhanced glucose clearance at 60 and 120 minutes post-challenge (Fig. 4H), and improved insulin sensitivity (Fig. 4I). To investigate a potential mechanism linking fermented food metabolites to glycemic improvement, we examined whether wGS-FDMs directly stimulate GLP-1 release from enteroendocrine cells. As accurate measurement of endogenous GLP-1 secretion in mice is notoriously unreliable with standard immonoassays^50,51^, we instead used the NCI-H716 line, a human colonic adenocarcinoma cell line ^52^, to directly assess wGS-FDM stimulation of GLP-1. Treatment with wGS-FDMs resulted in approximately 200–250% greater GLP-1 secretion relative to control media under 100 mM glucose conditions (Fig. 4J-K), demonstrating that wGS-FDMs can act directly on enteroendocrine cells to stimulate incretin release.

Together, these findings demonstrate that wGS-FDMs attenuate HFD-induced weight gain, remodel gut microbiota composition, reduce intestinal and systemic inflammatory infiltration, and improve glycemic control. Direct stimulation of enterendocrine GLP-1 secretion provides a candidate mechanism linking fermentation-derived metabolites modulate shared immune-metabolic axes to confer broad physiological benefit.

## Discussion

In this study, we leveraged a reverse translational strategy to investigate how vegetable-based fermented food (VBFF) metabolites shape host physiology, beginning with the clinical observation that fermented food consumption improves immune status in healthy adults ^29,53^. We found that in the absence of inflammatory challenge, wGS-FDMs reshape intestinal physiology, microbiota composition, and immune tone. Specifically, increased large intestinal length and elevated cecal CFUs reflect enhanced microbial fermentation capacity and gastrointestinal resilience, both associated with improved gut barrier integrity and reduced systemic inflammation in mouse models^54,55^. *Akkermansia muciniphila* was enriched by sauerkraut metabolites and has been linked to strengthened intestinal barrier function and increased longevity in both murine and human populations^56,57^. Expansion of TCRβ^+^ T cells and effector-skewed TCRβ^+^ T cells compartment suggest greater baseline T cell activation or homeostatic proliferation^58–60^. Concurrently, elevated CD80 expression on F4/80^hi^MHCII^lo^ macrophages suggests increased co-stimulatory capacity in this subset. However, given the observed attenuation of T cell responses upon challenge, this may instead reflect a toleragenic antigen presentation context, consistent with the known regulatory functions of this macrophage population^61–63^. Critically, the absence of immune remodeling in germ-free mice consuming wGS-FDM suggests that FDMs modulate the immune circuit in a microbiome dependent manner.

The mechanism by which wGS-FDMs engage the microbiome-immune axis remains to be fully defined. Future studies should determine to what extent different FDMs induce shifts in the composition or metabolic activity of the microbiome that may in turn calibrate host immune readiness. Alternatively, FDMs may act directly on the microbiota-matured immune system exerting activity that parallels the known ability of microbiota-derived metabolites (MDMs) such as short-chain fatty acids and secondary bile acids to shape T cell differentiation and macrophage function^10,64–67^. For example, the enrichment of aromatic amino acid derivatives, including 3-PLA and 4-HPLA, established ligands for the hydroxycarboxylic acid receptor 3 (HCA3) on immune cells, have been shown to modulate inflammatory signaling in macrophages^43,44^.

The wGS-FDM induced remodeling of immune tone we observed in a healthy mouse model raised the question of whether wGS-FDMs could be effective in improving outcomes in models of acute or chronic inflammation. In an acute inflammatory challenge model, wGS-FDM attenuated DSS-induced weight loss, severity of disease, and colitis across epithelial, immune and transcriptional readouts. Reduced *Ifng* expression points to restrained cytotoxic T cell activity, while elevated *Cldn3* and preserved colon length indicate that epithelial barrier integrity is maintained. Together, these findings support a model in which FDM-driven immune pre-conditioning selectively limits T cell-mediated epithelial damage rather than broadly suppressing inflammation, consistent with mechanisms previously described in the context of tumorigenesis^68^. wGS-FDM induced effector T cell enrichment and macrophage co-stimulatory priming reflect enhanced immune preparedness, calibrating the mucosal immune setpoint to constrain pathological escalation. The convergence of these findings with the *in vitro* NF-κB suppression data supports the notion that wGS-FDMs attenuate inflammatory severity.

In a high-fat diet model, wGS-FDM reduced weight gain and remodeled the microbiome through increased relative abundance of Lachnospiraceae family members and reduction of taxa associated with metabolic inflammation^69^ wGS-FDMs may partly exert their metabolic benefits through microbiome modulation, direct stimulation of GLP-1 secretion by wGS-FDMs in a cell culture model points to a mechanism by which wGS-FDMs could enhance insulin secretion and appetite regulation independent of the gut microbiome similar to that observed in a range of microbial metabolites and gut microbiota-derived secondary bile acids^70,71 72,73^. Additionally, among the metabolites identified during vegetable fermentation was N-lactoyl-phenylalanine, a recently described bioactive metabolite^46^ that increases following acute exercise and suppresses appetite. Similarly, branched chain hydroxyacids such as HMVA, which we observed in the wGS-FDMs, protect against HFD-induced obesity in mice^74^. The convergence of immune, microbiota, and endocrine data supports a model in which wGS-FDMs engage multiple parallel pathways to confer metabolic protection during HFD consumption.

Together, these findings position FDMs as potent host physiological regulators that confer measurable host benefits and operate through both microbiota-dependent and independent mechanisms. With the spectrum of metabolites that are inferred or known to bind to specific host receptors and elicit specific responses, an important question is whether the host physiological changes are due to single metabolites (or closely related classes of metabolites) acting through a definable small number of host pathways. At the other end of the spectrum of complexity, the effects on the host may be a result of a diverse array of FDMs acting on the host through multiple pathways, both directly and via microbiota metabolic modification. Additional investigations are needed to identify the specific bioactive metabolite or metabolites within wGS-FDMs responsible for the effects we observe. This work provides the foundation for investigating the molecular underpinnings of the health effects of fermented food consumption and positions FDMs as tractable targets for future therapeutics.

## Data Availability

Datasets and code for analysis are available at https://github.com/SonnenburgLab/MouseFeFo. Raw data files for metabolomics, 16S, and metagenomics sequencing available upon acceptance.

## Acknowledgments

We want to thank members of the Sonnenburg lab for discussions. In particular, Hollis Dupont, Tadashi Takeuchi, Tayler A. Sulse, as well as Steven Higginbottom and Cherelle Soriano. In addition, thank you to Miles Tyner, Megan Danielewicz, Catherine Liuo. Special thanks to, and to chef Andrew Mayne, chef Heidi Mitchell, and Stanford Dining Hospitality and Auxiliaries for providing support and space for the production of the fermented foods. Finally, thank you to Alice Kendall Duncombe Fessler.

## Author Contributions

E.B.C. designed, performed and analyzed all metabolomics experiments for all figures. ER.G. assisted with metabolomics sample preparation. E.K.R. and E.B.C. designed and performed all mouse experiments. J.F. helped with every mouse experiment and performed all flow cytometry experiments, E.K.R. analyzed all flow cytometry experiments for all figures. Y.G. performed germ free mouse experiments, while E.K.R. analyzed associated experiments. E.B.C. and A.R. designed, performed and analyzed ITT and GTT experiments. E.K.R. designed, performed, and analyzed all in vitro macrophage experiments. E.B.C. designed, performed and analyzed all in vitro GLP-1 experiments. E.K.R designed, performed and analyzed all RT-qPCR experiments. S.G.H. analyzed metagenomic data. E.B.C., E.K.R., E.D.S. and J.L.S. conceived and coordinated the project. E.B.C. and E.K.R. wrote the manuscript with input from all other coauthors. S.P.S., E.D.S. and J.L.S. provided funding.

## Ethics declaration

The authors declare no relevant competing interests.

## Supplementary Information

Supplementary Table 1 - Metabolomics Sample List

Supplementary Table 2 - Metabolomics Results

Supplementary Table 3 - 16S Results

Supplementary Table 4 - Metagenomics Results

Supplementary Table 5 - Antibody List

## Materials and Methods

### Ethics statement

All mouse experiments were approved by the Stanford University Institutional Animal Care and Use Committee (IACUC) and conducted in accordance with institutional guidelines for the humane care and use of laboratory animals.

### Fermented food production and sampling

Vegetable fermentations were carried out under spontaneous lactic acid fermentation conditions in sterile glass bottles. For green sauerkraut and red sauerkraut, common or red cabbage was sliced into thin ribbons after removal of the core. A total of 2% NaCl (w/w) was added, and the salted cabbage was mixed and allowed to sit for one hour with periodic massaging to extract natural brine. Extracted brine was reserved, and salted cabbage was packed into sterile jars; remaining brine was distributed evenly across jars to fully submerge the cabbage. For the starter-inoculated condition (Green Sauerkraut with starter; sGS), 5% (w/w) of commercial sauerkraut brine from a late-stage ferment was added to freshly prepared jars to enrich for taxa associated with the terminal stages of fermentation. Curtido was prepared similarly, with the addition of carrots, onions, and oregano. Baechu kimchi was prepared with napa cabbage combined with a paste of garlic, Korean red chili pepper, salt, water, and sugar, with a subset of preparations including fish sauce. Nabak kimchi was prepared as a water kimchi with a brine of scallions, garlic, ginger, radish, carrots, and Korean pear. Samples were collected at multiple timepoints spanning days 0, 4, 7, 21, and 60, immediately frozen on dry ice, and stored at −80°C until analysis. Final timepoints were determined based on pH stabilization.

### Fermented food extraction and preparation for *in vitro* and *in vivo* assays

For metabolite extraction, 100 µL of fermented food brine was added to 1ml of HPLC-grade methanol and incubated at −20°C for 1 hour. Samples were centrifuged at 20,000 × g for 10 min and the supernatant was transferred and dried under a stream of nitrogen using a TurboVap. For *in vitro* assays, dried extracts were resuspended in 100 µL of HPLC-grade water. Metabolite extracts were used at a 1:100 dilution in cell culture medium unless otherwise specified. For *in vivo* gavage experiments, dried extracts were resuspended in 100 µL of mouse facility di-water, with 300 µL of extract was administered per animal per day. Unfermented cabbage metabolite extract (CB-M) was prepared from equivalent quantities of fresh, salted cabbage subjected to the same extraction protocol and served as a non-fermented substrate control.

### Semi-targeted and targeted metabolomics

Semi-targeted LC-MS was performed as previously described^1^. Briefly, 100 µL of brine was added to 1ml of HPLC-grade methanol containing internal standards and incubated at room temperature for 5 min, followed by centrifugation at 5,000 × g for 10 min. Supernatants were dried and reconstituted in 50:50 methanol:water (v/v) with internal standards. Samples were analyzed on an Agilent Q-TOF 6545 mass spectrometer using C18 in positive and negative mode. Compound annotation was performed using MS-DIAL (v3.82) with an authentic standard reference library and a custom gut microbiome-focused database. The area under the curve for each metabolite was normalized to the sum of internal standard signals per sample. For metabolites detected across multiple acquisition modes, the normalized peak area from the mode with the highest signal-to-noise ratio was used. For targeted metabolomics, a custom database was additionally used for annotation of metabolites of interest. Pairwise Jaccard indices were calculated for significantly changing metabolites (two-sided Wilcoxon rank-sum test, FDR-adjusted p < 0.05, |log2FC| > 1) to quantify compositional similarity across ferments.

### Animal models

#### Healthy conventional microbiota mouse model

Seven-week-old male C57BL/6J mice (Jackson Laboratory) were randomized to daily oral gavage with sauerkraut fermentation derived metabolites (SK-FDMs) (300 µL) or sterile water for 17 days. Body weight was recorded daily. At endpoint, the small intestine, large intestine, spleen, and cecum were collected for length and weight measurements, and tissues were processed for flow cytometry, gene expression analysis, and epithelial proliferation assays as described below.

#### DSS-induced colitis model

Eight-week-old conventional male C57BL/6J mice (Jackson Laboratory) were gavaged daily with either SK-FDMs or water for seven weeks prior to colitis induction. Colitis was then induced by providing 2.5% (w/v) dextran sulfate sodium (DSS; 36–50 kDa, MP Biomedicals) in autoclaved drinking water for seven days, followed by a three-day recovery phase on autoclaved water. Daily gavage of FDMs was continued throughout DSS exposure and recovery. Body weight was recorded daily; Disease Activity Index (DAI) was calculated as a composite score of weight loss, stool consistency, and fecal occult blood. Body weight loss was scored as 0 (none), 1 (1–5%), 2 (5–10%), 3 (10–20%), or 4 (>20%); stool consistency as 0 (normal), 2 (loose stool), or 4 (diarrhea); and stool blood as 0 (negative), 2 (fecal occult blood positive), or 4 (gross bleeding)^2^.

At experimental endpoint, colon length, cecal weight, and immune cell populations were assessed. Total CD45+ cell counts were measured by flow cytometry of large intestinal lamina propria preparations. Gene expression in colonic tissue was assessed by RT-qPCR.

#### High-fat diet model

Conventional-microbiome male C57BL/6J mice (Jackson Laboratory) were placed on a 60% kcal fat diet (Research Diets D12492) for one week prior to treatment initiation. Mice were then randomized to receive water, CB-M, or SK-FDMs by oral gavage of 300ul daily for nine weeks while maintained on the HFD. Body weight was measured weekly. Stool was collected for 16S rRNA amplicon sequencing at weeks 0, 5, 7, and 11. Fasting blood glucose was measured after a 6-hour fast using a OneTouch glucometer. Glucose tolerance tests (GTTs) were performed by intraperitoneal injection of 2 g/kg glucose, and insulin tolerance tests (ITTs) were performed with 0.75 U/kg insulin i.p., with blood glucose measured at 0, 15, 30, 60, 90, and 120 min post-injection.

#### Germ-free mouse model

Gnotobiotic germ-free C57BL/6J mice were housed in flexible film isolators at the Stanford Gnotobiotic Facility^3^. Mice were supplied with filter-sterilized wGS-FDM (concentration) in their sterile water or sterile water for 17 days. At the endpoint, intestinal tissues were collected for length measurements, histological analysis, epithelial proliferation, and immune profiling as described above. All handling was performed under sterile conditions to maintain germ-free status throughout.

### Intestinal immune cell isolation and flow cytometry

Single-cell suspensions from the small intestine (SI), large intestine (LI), and spleen were prepared as previously described ^4^ with minor modifications. Mice were euthanized by CO_2_ inhalation, and tissues were rapidly dissected. Intestinal tissues were cleaned of mesenteric fat, Peyer’s patches were removed, and luminal contents were flushed with ice-cold RPMI. Tissues were opened longitudinally, rinsed, and cut into 1–2 cm segments before transfer into strip buffer (RPMI containing 5 mM EDTA and 1 mM DTT). Samples were shaken at 220–250 rpm for 20 min at 37°C, and supernatants were collected through fine-mesh strainers to the intraepithelial lymphocyte (IEL) fraction. Two additional EDTA stripping steps were performed to ensure complete epithelial removal. Remaining tissue fragments were minced and transferred to digestion buffer (RPMI supplemented with 1 mg/mL Liberase TM and 0.5 mg/mL DNase I) and incubated for 25 min at 37°C with gentle agitation to release lamina propria (LP) leukocytes. Digestion was quenched with fetal bovine serum (FBS), and resulting suspensions were passed sequentially through 70-µm and 40-µm strainers, washed with RPMI 1640 + 2% FBS, and pelleted by centrifugation (500 × g, 5 min). Cell pellets were resuspended in complete RPMI (RPMI + 2% FBS) prior to staining.

Single-cell suspensions were stained with fluorophore-conjugated antibodies in FACS buffer (PBS + 2% FBS + 2 mM EDTA) for 30 min at 4°C (see Supplemental Table 5 for a list of Abs, clones, fluorochromes, and manufacturers). Live/dead discrimination was performed using LIVE/DEAD Fixable Aqua or Near-IR (Thermo Fisher). Intracellular staining for Foxp3 and Ki67 was performed using the Foxp3/Transcription Factor Staining Buffer Set (eBioscience). Data was acquired on a Cytek Aurora and analyzed using FlowJo v10. Gating strategies used FSC/SSC for singlet discrimination followed by live cell gating. Immune populations were identified using standard surface marker combinations. TCRβ+ T cells, CD4+ and CD8+ subsets, F4/80hiMHCIIhi activated macrophages, marginal zone macrophages, and resting macrophages were identified within the CD45+ gate. EpCAM+ cells were gated within the CD45− population. For splenic macrophage activation, CD80 MFI was used as a marker of antigen-presenting capacity. Cell counts were normalized to counting beads.

### NF-κB reporter macrophage assay

The murine NF-κB-GFP reporter macrophage line used in this study was generated as previously described ^5^. Briefly, a bone marrow-derived cell line stably expressing GFP under the control of five tandem NF-κB response elements upstream of a minimal CMV promoter (5× NF-κB-CMVmin-GFP) was used as the primary murine reporter system. Cells were maintained in Dulbecco’s Modified Eagle Medium (DMEM) supplemented with 10% heat-inactivated fetal bovine serum (FBS), 2 mM L-glutamine, and 1% penicillin-streptomycin at 37°C in 5% CO_2_.

For the assay, cells were seeded in 96-well plates at 1 × 10^5^ cells per well 24 hours prior to treatment. Cells were pre-treated for 20 hours with SK-FDMs, CB-M, or vehicle control (HPLC-grade water) at a 1:100 dilution in complete DMEM. Following pre-treatment, cells were stimulated with lipopolysaccharide (LPS) (200 ng/mL; Sigma L2630) for 18 hours. GFP fluorescence intensity was quantified by flow cytometry on a Acea Novocyte Quanteon. Cells were gated using (forward scatter-area) FSC-A and (side scatter-are) SSC-A to select live cells, and then gated for singlets using FSC-A/FSC-H prior to GFP median fluorescence intensity (MFI) quantification. Data were analyzed in FlowJo v10, and GFP MFI was normalized to the vehicle-stimulated control condition. Cell viability was assessed in parallel using LIVE/DEAD Fixable Aqua staining; no significant reduction in viability was observed across any treatment condition.

For human macrophage experiments, THP-1 monocyte-derived cells were used. The THP-1 NF-κB-GFP reporter line was generated as previously described^6^. Cells were cultured in RPMI 1640 supplemented with 10% low-endotoxin FBS and 1× penicillin-streptomycin, and incubated at 37°C in 5% CO2. Cells were seeded at 1 × 10^5^ cells per well in 96-well plates, pre-treated with SK-FDMs or controls for 2 hours, and then stimulated with LPS (200 ng/mL) for 18 hours. GFP MFI was quantified by flow cytometry and normalized to vehicle controls as described above. Assay validity was confirmed by ensuring that the MFI of positive (LPS-stimulated) controls exceeded unstimulated controls by at least 10-fold.

### Cytokine quantification

TNF-α and IL-6 protein concentrations in macrophage supernatants were measured by ELISA (R&D Systems) following the manufacturer’s instructions. Briefly, macrophages were seeded at 1 × 10^5^ cells per well in 96-well plates, pre-treated with SK-FDMs, CB-M, or vehicle for 2 hours, and stimulated with LPS (200 ng/mL) or left unstimulated for 18 hours. Supernatants were collected by centrifugation (500 × g, 5 min) and stored at −80°C until analysis. Absorbance was measured at 450 nm on a SpectraMax plate reader.

### NCI-H716 enteroendocrine cell GLP-1 secretion assay

Human enteroendocrine NCI-H716 cells (ATCC CRL-3281) were maintained in suspension culture in RPMI-1640 supplemented with 10% FBS, 2 mM L-glutamine, and 1% penicillin-streptomycin at 37°C in 5% CO_2_. For secretion assays, cells were counted by hemocytometry, centrifuged at 300 × g for 5 min, and resuspended in serum-free RPMI. Cells were plated at 5 × 10^5^ cells per well in Matrigel-coated 96-well plates and allowed to equilibrate for 48 hours prior to stimulation. A Dipeptidyl peptidase-4 (DPP-IV) inhibitor (Millipore, final concentration 10 µM) was included in all assay buffers to prevent GLP-1 degradation. Each condition was assayed in technical triplicate. Cells were incubated with SK-FDMs (1:100), CB-M (1:100), or vehicle in serum-free RPMI supplemented with 100 mM glucose for 2 hours. Plates were centrifuged at 300 × g for 5 min and supernatants were collected and stored at −80°C until analysis. Total GLP-1 was quantified using the Meso Scale Discovery (MSD) Total GLP-1 assay kit (K1514PK), following the manufacturer’s instructions.

### RNA extraction and quantitative RT-PCR

Colonic or intestinal tissue (30–50 mg) was homogenized in RLT buffer (Qiagen), and total RNA was extracted using the RNeasy Mini Kit (Qiagen) following the manufacturer’s protocol, including an on-column DNase I digestion step. RNA concentration and purity were assessed by Nanodrop spectrophotometry. cDNA synthesis was performed from 500 ng of total RNA using the iScript cDNA Synthesis Kit (Bio-Rad). Quantitative PCR was performed using iTaq Universal SYBR Green Supermix (Bio-Rad) on a CFX96 Real-Time PCR System (Bio-Rad) with the following cycling conditions: 95°C for 3 min, followed by 40 cycles of 95°C for 10 s and 60°C for 30 s. Primer sequences for all target genes in table below.

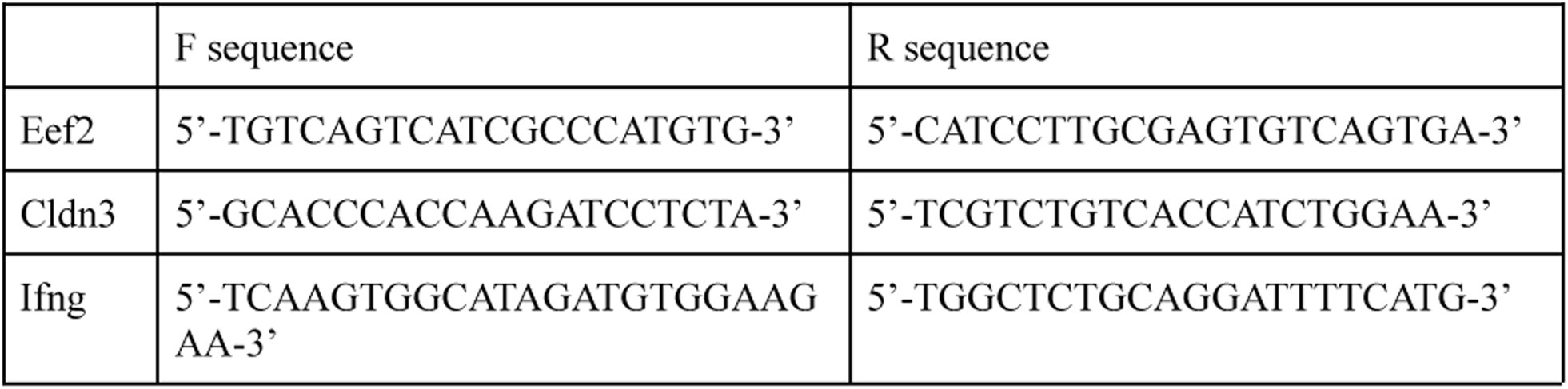

Gene expression was normalized to the housekeeping gene Eef2 using the ΔΔCt method. Genes analyzed included Ifng and Cldn3.

### Sequencing

#### 16S rRNA microbiota sequencing and analysis

Cecal or stool contents were collected into sterile microcentrifuge tubes and immediately frozen at −80°C. DNA extraction was performed using the DNeasy PowerSoil Pro Kit (Qiagen) following the manufacturer’s instructions. The V4 region of the 16S rRNA gene was amplified using primers 515F/806R and sequenced on an Illumina MiSeq platform (2 × 250 bp paired-end). Amplicon sequence variants (ASVs) were resolved using DADA2 (v1.18) with standard quality filtering, denoising, and chimera removal. Taxonomic assignment was performed against the SILVA database (v138). Alpha and beta diversity analyses were performed in R using the vegan (v2.7-3) packages. Community-level differences between groups were assessed by PERMANOVA using Bray-Curtis dissimilarity (999 permutations). Differential abundance analysis was performed using DESeq2 with Benjamini-Hochberg FDR correction. Receiver-operating characteristic (ROC) analysis with 100% AUC was used to identify ASVs maximally discriminating between diet groups.

### Metagenomic sequencing and analysis

Fecal matter was submitted to the University of Wisconsin-Madison Biotechnology Center. DNA was isolated from 250mg of fecal matter using the DNeasy 96 PowerSoil Pro QIAcube HT Kit (QIAGEN, Hilden, Germany).DNA concentration was verified using the Quant-iT™ PicoGreen® dsDNA Assay Kit (ThermoFisher Scientific, Waltham, MA, USA). Samples were prepared according to the QIAGEN FX DNA Library Preparation Kit (QIAGEN). Quality and quantity of the finished libraries were assessed using an Agilent Tapestation (Agilent, Santa Clara, CA) and Qubit® dsDNA HS Assay Kit, respectively. Paired end, 150 bp sequencing was performed using the Illumina NovaSeq X Plus (Illumina, San Diego, CA).

#### Preprocessing and taxonomic classification

Raw reads were processed using the nf-core/mag pipeline (https://dx.doi.org/10.1038/s41587-020-0439-x; 10.1093/nargab/lqac007) with MEGAHit selected as the assembler. Taxonomy was assigned with the nf-core.taxprofiler (https://dx.doi.org/10.1038/s41587-020-0439-x; 10.1101/2023.10.20.563221) pipeline using MetaPhlAn4 for taxonomic classification with the CHOCOPhlAn vJan25 database (https://doi.org/10.1038/s41587-023-01688-w). Downstream analyses were performed in R (v4.5.2).Within both species- and genus-level taxonomic levels, “unclassified” ranks were removed. The classifications were then re-normalized to sum to 100% so that bars reflect the proportion of classified reads at that level (Supplementary Fig. 7A). Mean relative abundance was calculated per Treatment x Diet Group. Taxa with an overall sample mean below 1% were collapsed into a single “Low Abundance” category. Stacked bar charts were visualized with ggplot2 (v4.0.2).

#### Beta diversity

Bray-Curtis dissimilarity was computed on the species- and genus-level relative abundance tables using the “vegdist” function in vegan (v2.7.3). Principal Coordinates Analysis (PCoA) (Supplementary Fig 14A) was performed with the “cmdscale” function in vegan (v2.7.3). To assess treatment effects in the context of host diet, two complementary ordination strategies were applied for the Water and 3-week Fermented Sauerkraut treatment groups: (1) a combined ordination in which all samples were included in a single distance matrix and treatment groups were included in a single distance matrix and diet groups were shown as separate facets, and (2) separate per-diet ordinations in which each diet’s samples were independently ordinated to isolate within-diet treatment variation. Ellipses were drawn per Treatment x Diet group. Visualization was done with ggplot2 (v4.0.2).

#### Differential abundance

Differentially abundant taxa were identified using “ANCOMBC2” from the ANCOMBC (v2.12.1) package. Analyses were conducted separately for each diet at both the species and genus level. Because ANCOM-BC2 requires count data, MetaPhlAn relative abundances were converted to pseudo-counts by multiplying each sample’s relative abundance values by its post quality-controlled read count as determined from the MultiQC (10.1093/bioinformatics/btw354) summary statistics included in nf-core/taxprofiler (https://dx.doi.org/10.1038/s41587-020-0439-x; 10.1101/2023.10.20.563221) and then rounding to the nearest integer. ANCOM-BC2 was run with Treatment as a fixed effect. P-values were adjusted for multiple comparisons with the Benjimini-Hochberg (BH) method. Taxa were considered significant at a BH-adjusted p < 0.05. Results were visualized with volcano plots generated with ggplot2 (v4.0.2).

To assess whether sauerkraut consumption counteracts HFD-induced gut microbiome dysbiosis, we performed a rescue correlation analysis. First, a “dysbiosis score” was estimated for each taxon by running ANCOM-BC2 on water-only samples containing both conventional and high-fat diet mice, yielding a log2fold change reflecting the magnitude and direction of HFD-induced compositional shifts (positive = enriched in HFD, negative = depleted). Second, a “rescue score” was estimated by running ANCOM-BC2 on HFD samples only comparing 3w fermented to water, yielding a log2 fold change reflecting the effect of sauerkraut within the HFD context. Both models used read counts calculated as above. Taxa were then classified into four quadrants based on the direction of both scores. The correlation between dysbiosis and rescue scores was assessed using Spearman correlation, and an ordinary least squares regression line was overlaid alongside a theoretical “perfect rescue” line (slope = -1, intercept = 0), where points on this line would indicate that sauerkraut exactly counteracts the HFD effect.

### Statistical analysis

All statistical analyses were performed in R (v4.3) or GraphPad Prism (v10). For comparisons between two groups, two-tailed Mann-Whitney U tests were used for non-normally distributed data, and Student’s t-tests were applied when normality was confirmed by Shapiro-Wilk testing. For comparisons across three or more groups, one-way ANOVA with Tukey’s post hoc test was used. Longitudinal body weight and DAI data were analyzed by two-way ANOVA with Sidak’s multiple comparisons correction. Microbiota community differences were assessed by PERMANOVA (adonis2, vegan package) using Bray-Curtis dissimilarity with 999 permutations. Differential metabolite abundance was assessed using two-tailed Wilcoxon rank-sum tests with Benjamini-Hochberg FDR correction (adjusted p < 0.05). Statistical significance thresholds: * p < 0.05, ** p < 0.01, *** p < 0.001, **** p < 0.0001. All graphs show individual data points overlaid on box plots (median, IQR, 1.5× IQR whiskers) unless otherwise indicated. Sample sizes for each experiment are indicated in the corresponding figure legends.

---

**Supplementary Figure 1.**
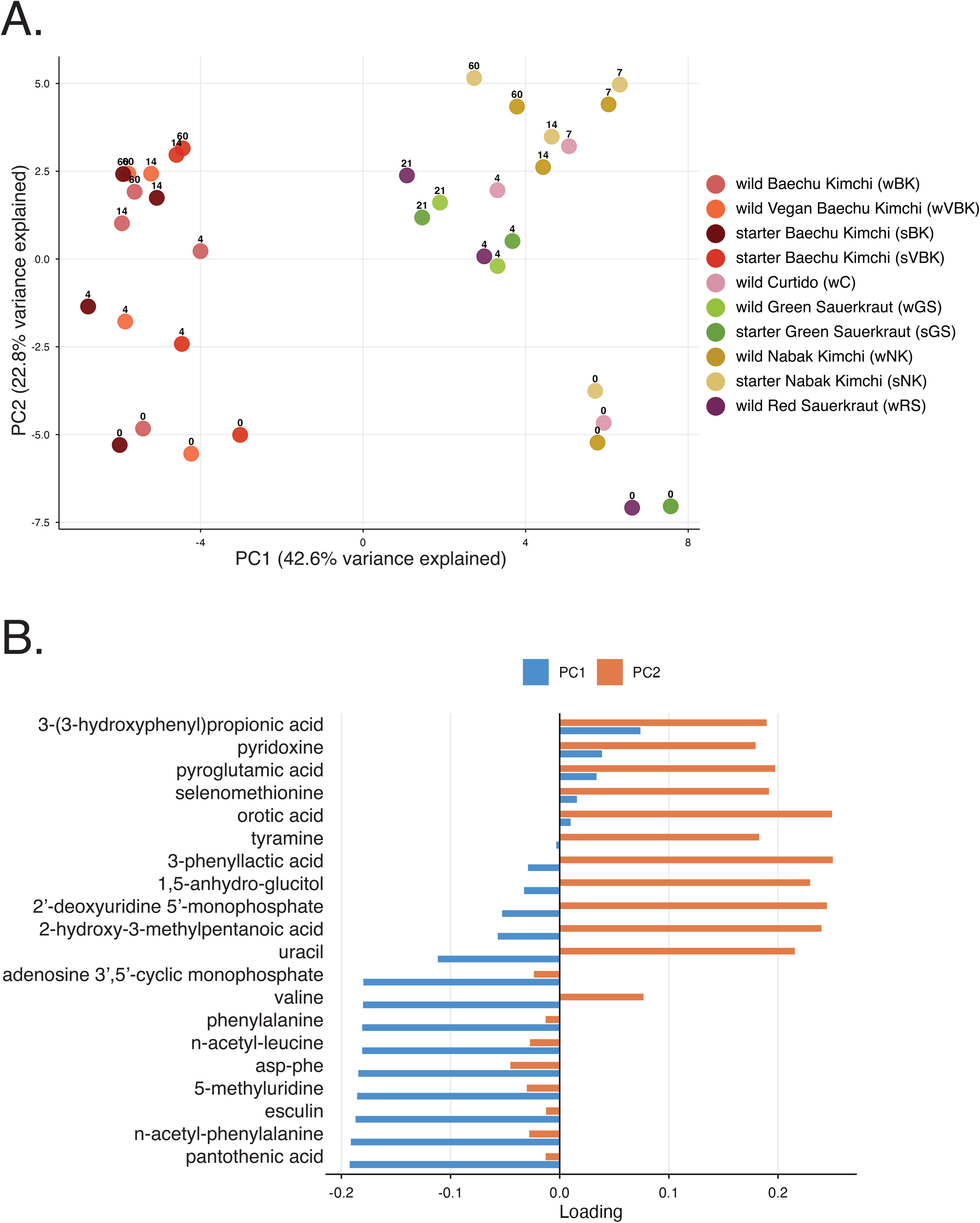
Metabolomics characterization of vegetable based fermented foods (VBFF). (A) Probabilistic principal component analysis (PPCA) of untargeted metabolomics profiles across all fermented food samples and fermentation stages. Each point represents a biological replicate; colors indicate food and fermentation conditions. PC1 and PC2 explain 42.6% and 22.9% of total variance, respectively. (B) PCA loadings for PC1 and PC2 showing the top metabolites driving sample separation.

**Supplementary Figure 2.**
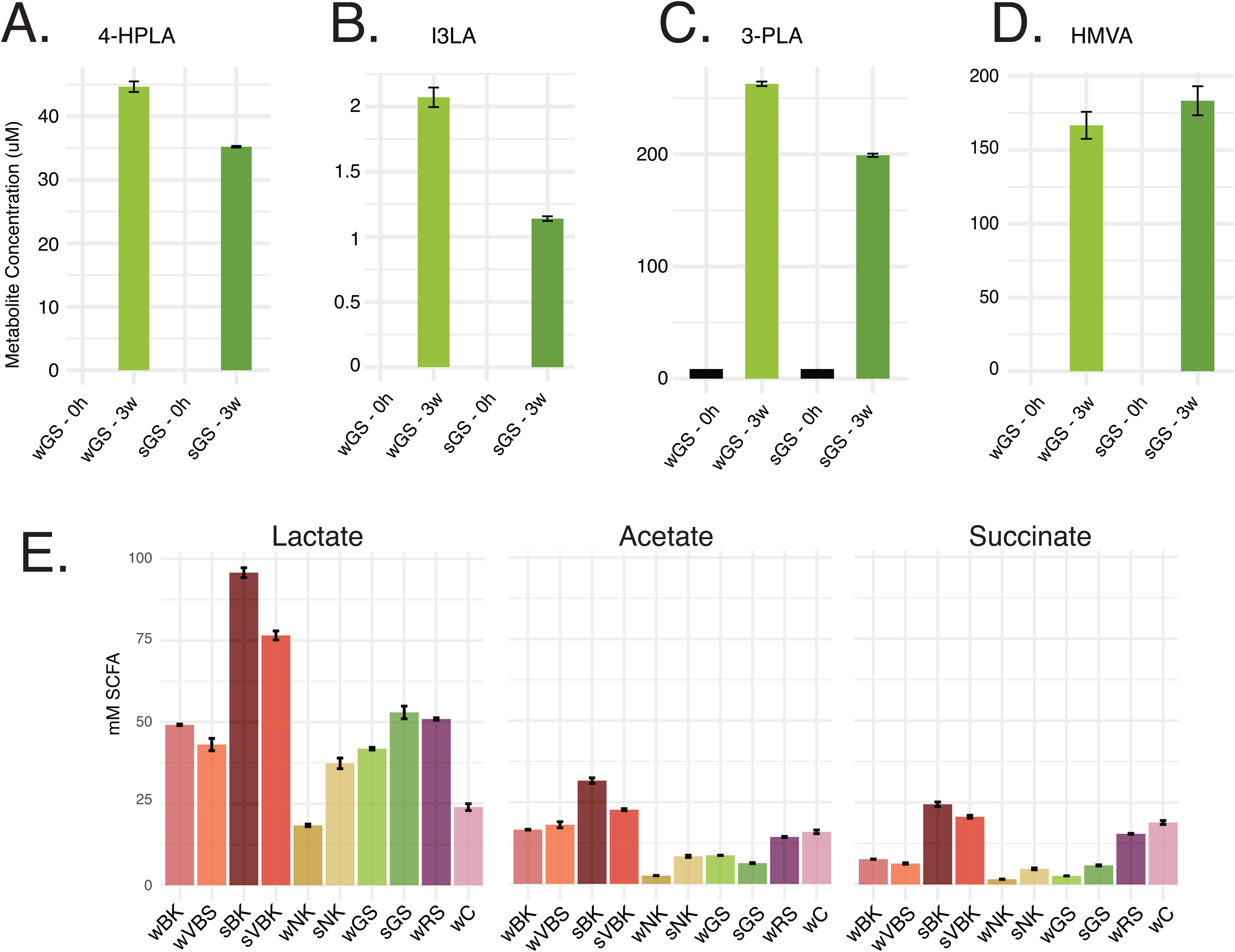
Targeted quantification of metabolites of interest and short-chain fatty acids (SCFAs). (A) Targeted LC-MS/MS quantification of 4-hydroxyphenyllactic acid (4-HPLA), (B) indole-3-lactic acid (I3LA), (C) 3-phenyllactic acid (3-PLA), (D) and HMVA (µM) in green sauerkraut fermented with (sGS) or without starter culture (wGS) at 0 hours and 3 weeks, validating untargeted metabolomics findings. Values represent mean ± SEM. (See Supplementary Table 2.) (E) SCFA quantification (mM) of lactate, acetate, and succinate across all fermented food conditions at the final fermentation timepoint (3 weeks), confirming active lactic acid fermentation. Values represent mean ± SEM. (See Supplementary Table 2.)

**Supplementary Figure 3.**
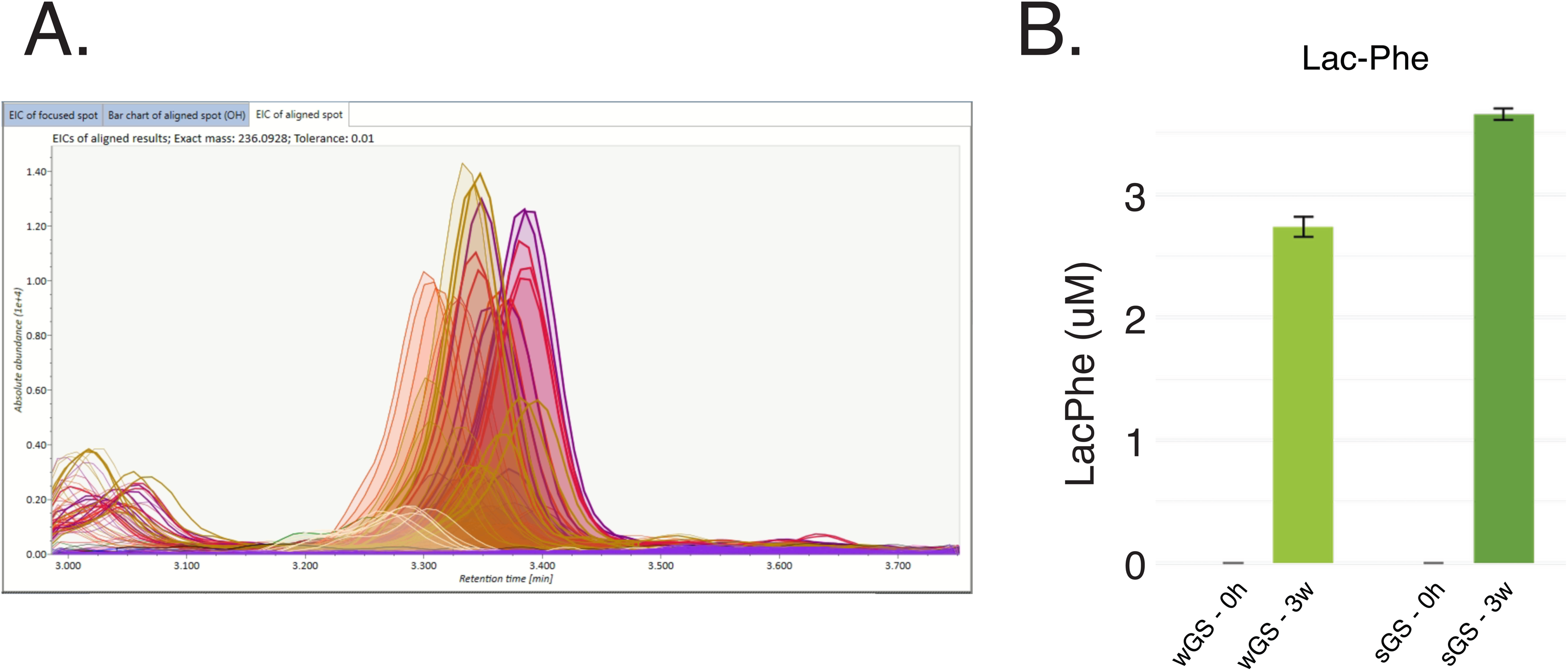
Identification and targeted quantification of Lac-Phe. (A) Extracted ion chromatograms from MS-DIAL of mass-to-charge ratio (m/z) corresponding to N-lactoyl-phenylalanine (Lac-Phe) (m/z = 236.0928, negative mode) across all fermented food samples at multiple fermentation stages. (B) Lac-Phe concentrations measured by targeted LC-MS/MS in Green Sauerkraut fermented with (sGS) or without starter culture (wGS) at 0 hours and 3 weeks. Lac-Phe was quantified using an authentic synthetic standard and confirmed by matching retention time and Multiple Reaction Monitoring (MRM) transition (236.1→147 m/z, 236.1→88.1 m/z). Values represent mean ± SEM.

**Supplementary Figure 4.**
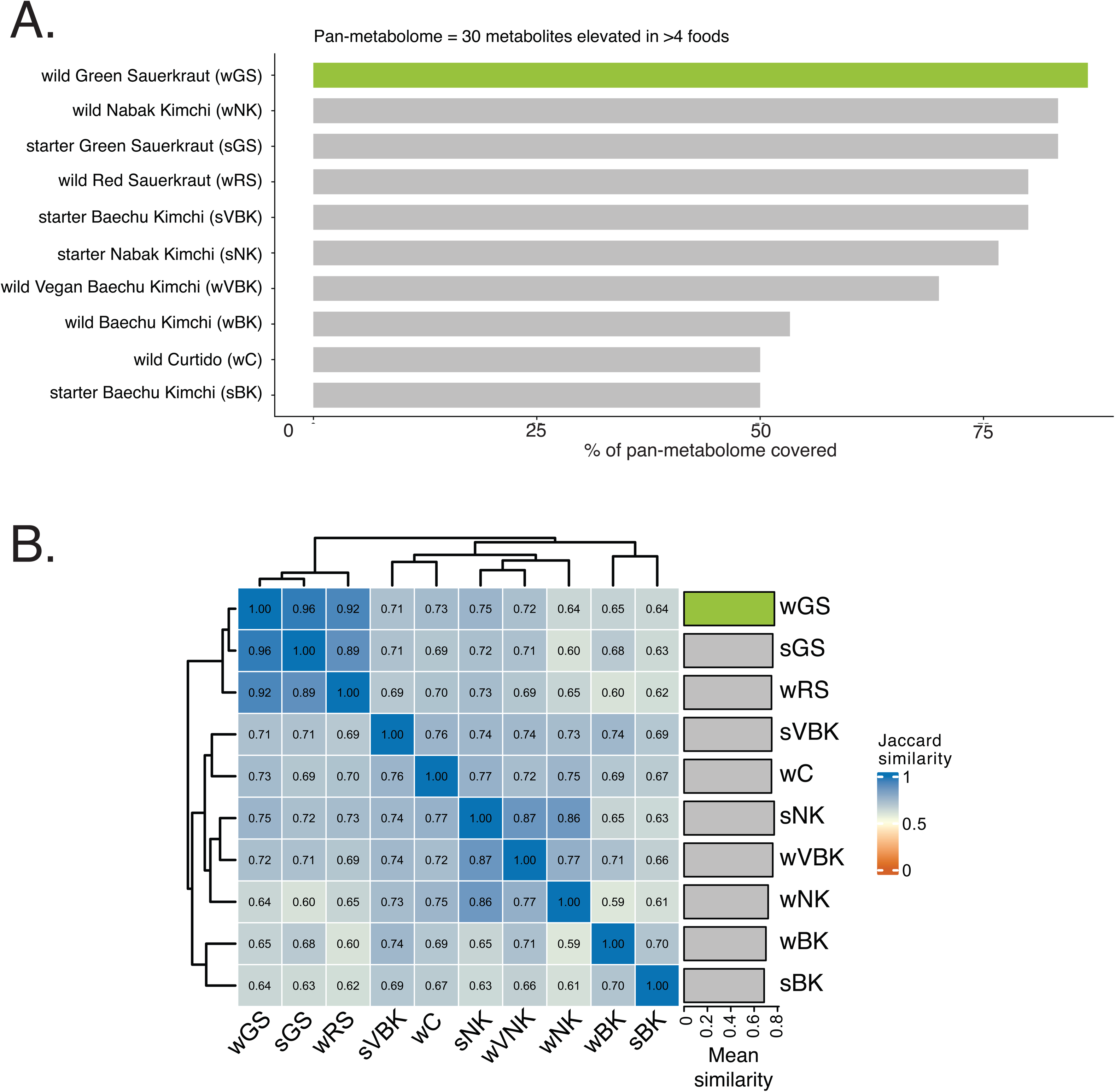
Pan-metabolome coverage. (A) The pan-fermented-food metabolome was defined as metabolites enriched (log₂FC > 2) in at least 4 fermented foods. Bars show the percentage of pan-metabolome metabolites covered by each condition. Green Sauerkraut (wGS; highlighted) showed the highest coverage and was selected as the primary food model. (B) Jaccard similarity heatmap across all fermented food conditions based on shared upregulated metabolites. Values represent pairwise similarity scores (0 = no overlap, 1 = identical); clustering was performed using Euclidean distance. Bar annotation (right) shows mean similarity per food across all others.

**Supplemental Figure 5.**
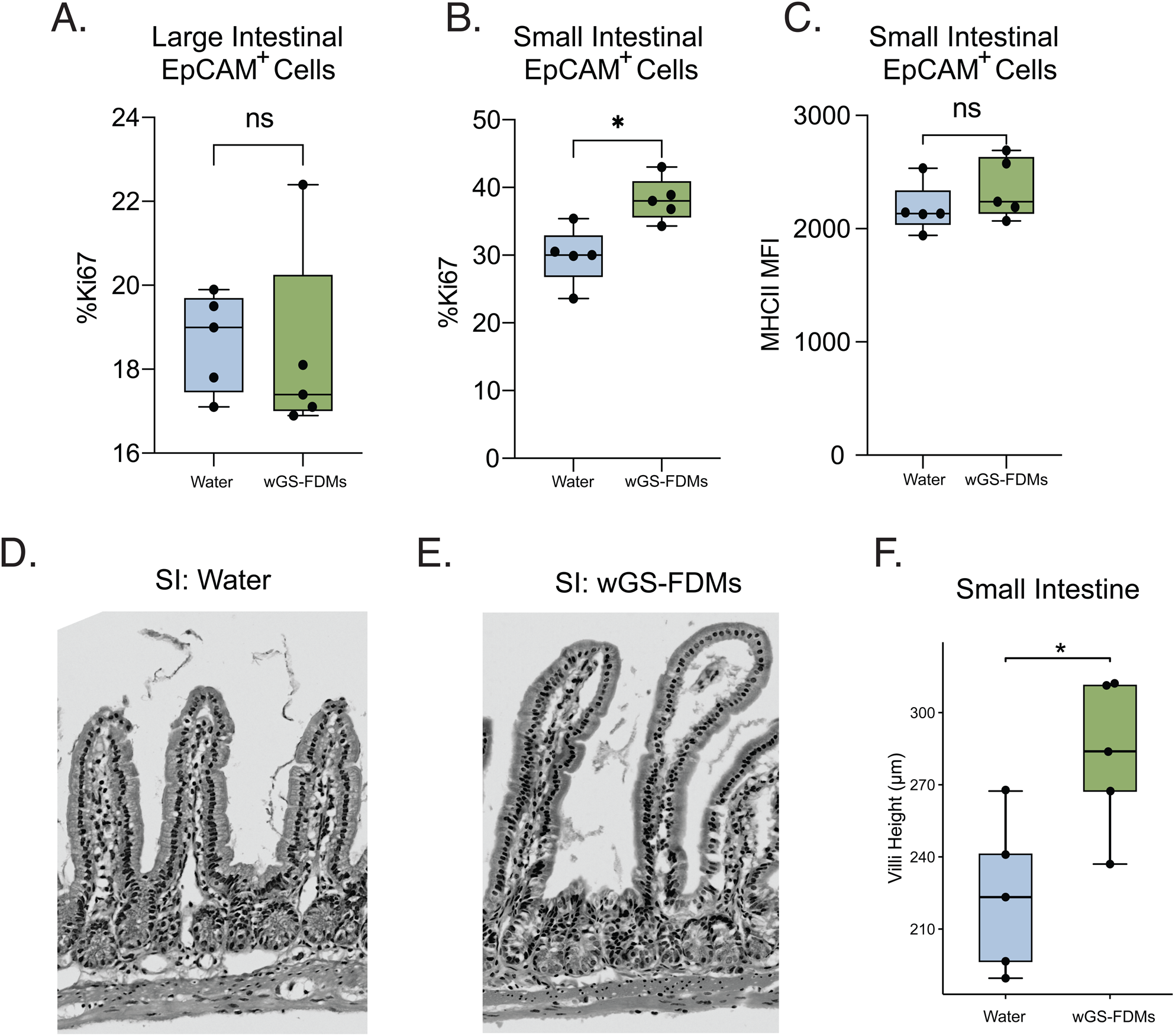
wGS-FDMs promote small intestinal epithelial growth without altering immune activation. (A) Percent of Ki67⁺ proliferating cells among EpCAM⁺ in large intestinal or (B) small intestinal epithelial cells for water and wGS-FDM-treated mice. wGS-FDM treatment was associated with increased epithelial proliferation in the small intestine, but not in the large intestine. (C) Major histocompatibility complex class II (MHCII) expression (mean fluorescence intensity (MFI)) on EpCAM⁺ small intestinal epithelial cells from water and wGS-FDM-treated mice. (D–E) Representative hematoxylin and eosin (H&E)-stained sections of the small intestine from water-treated and wGS-FDM-treated mice, showing villus morphology. (F) Quantification of villus height (μm) in the small intestine of water and wGS-FDM-treated mice. Data are presented as box-and-whisker plots with individual data points overlaid. Statistical significance was determined by Mann-Whitney U test or unpaired Student’s t-test. *p < 0.05; ns, not significant. n = 5–7 mice per group.

**Supplemental Figure 6.**
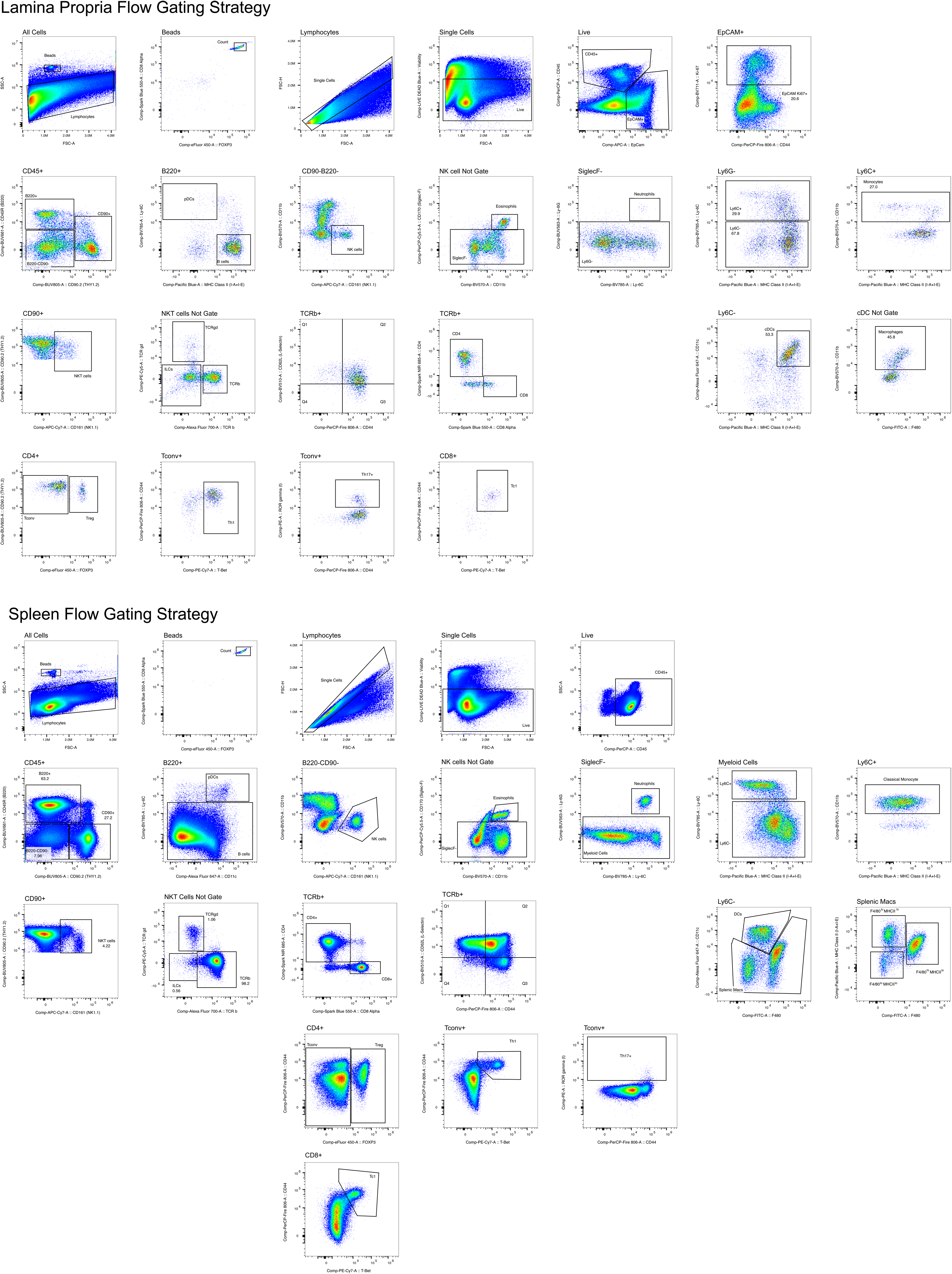
Gating strategy for spleen and lamina propria. Representative gating plots showing the sequential gating hierarchy used to identify immune populations in the small intestinal lamina propria (SI-LP), large intestinal lamina propria (LI-LP), and spleen.

**Supplemental Figure 7.**
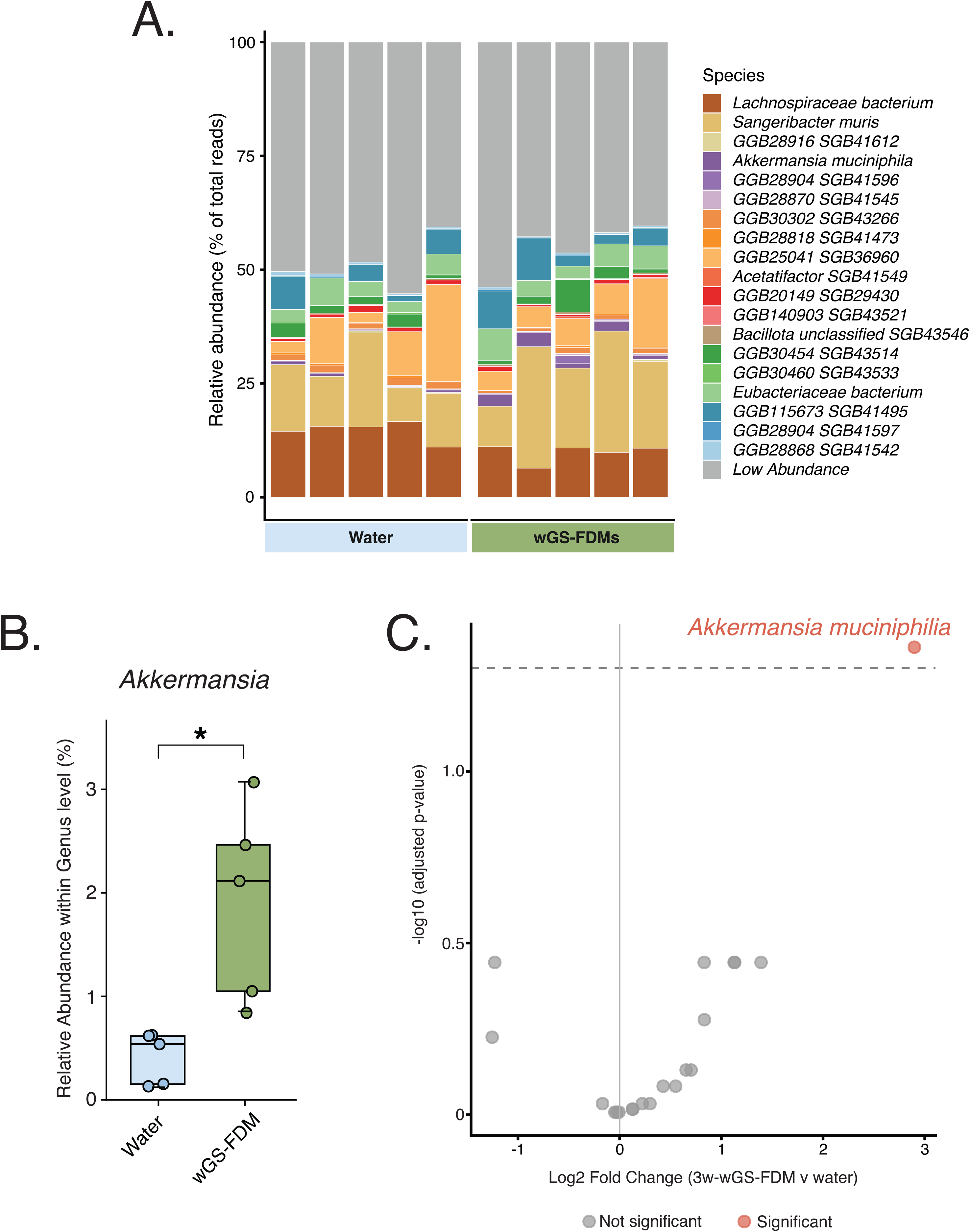
Impact of wGS-FDMs on microbiome composition and *Akkermansia* relative abundance. (A) Relative abundance of microbial species in cecal content of water and wGS-FDM-treated mice (n = 5 per group), with taxa contributing less than a defined threshold (<1%) grouped as ’Low Abundance’. (B) Genus-level Mann-Whitney U test for genus *Akkermansia* (BH-adjusted p-value). (C) Differential abundance comparing mice treated with water or 3w-wGS-FDMs (BH-adjusted).

**Supplemental Figure 8.**
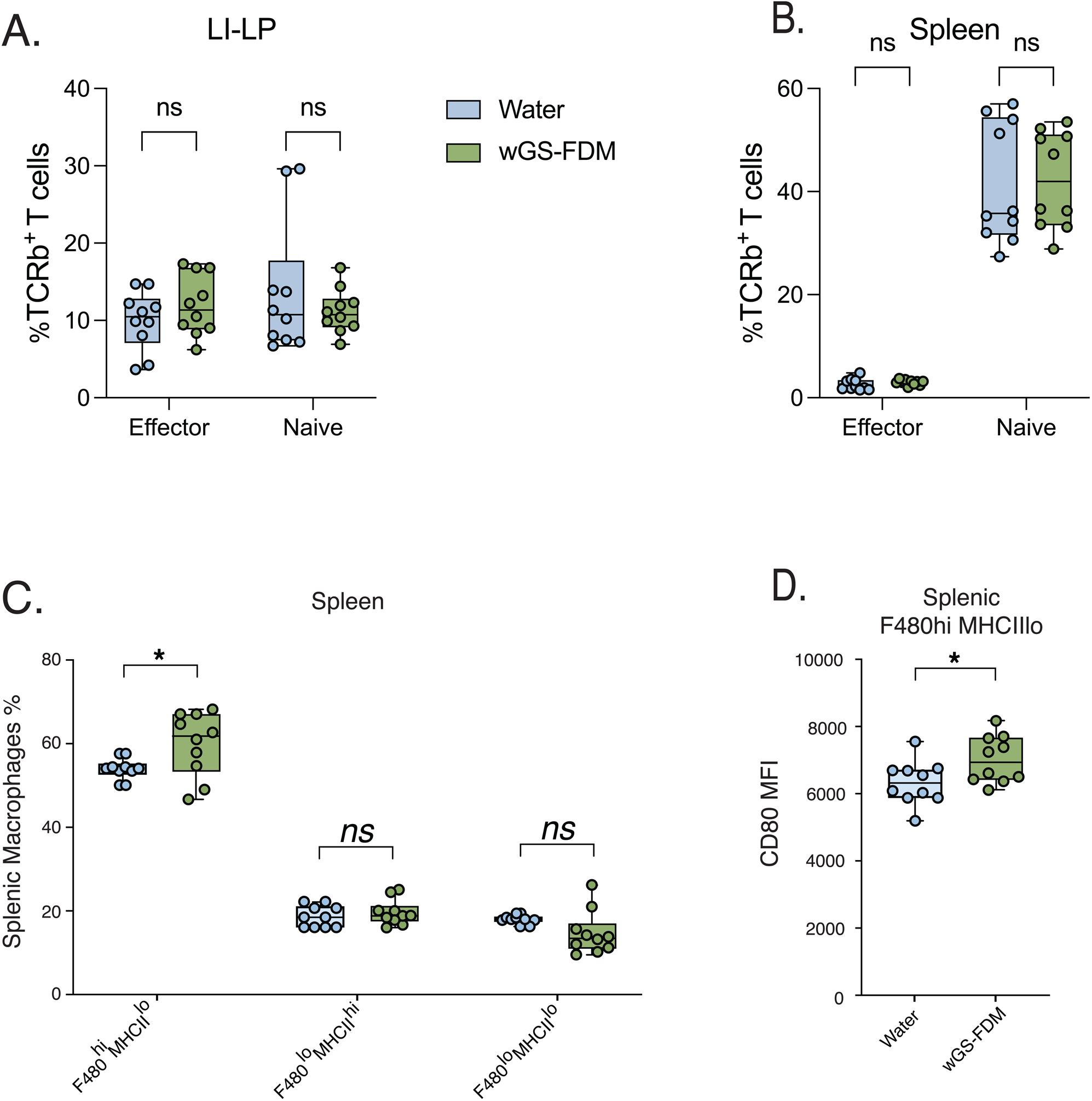
wGS-FDM treatment does not affect T cell proportions but increases splenic F480^hi^MHCII^lo^ macrophage frequency and CD80 expression. (A–B) Frequency of TCRβ⁺ T cells among total live cells in (A) the large intestinal lamina propria (LI-LP) and (B) spleen, gated on effector and naïve T cell populations. (C) Frequency of splenic macrophage subsets — F480^hi^MHCII^lo^, F480^lo^MHCII^hi^, and F480^lo^MHCII^lo^ — as a percentage of total splenic macrophages in water- and 3w SK-treated mice. (D) CD80 mean fluorescence intensity (MFI) on splenic F4/80^hi^MHCII^lo^ macrophages from water- and wGS-FDM-treated mice. Mice received either water (blue) or 3-week SK treatment (3w SK, green). Each dot represents an individual mouse. Boxes indicate interquartile range with median; whiskers show min/max values. *, p < 0.05; ns, not significant (unpaired t-test or Mann-Whitney test). Data are pooled from 2 independent experiments with n = 10 mice per group.

**Supplemental Figure 9.**
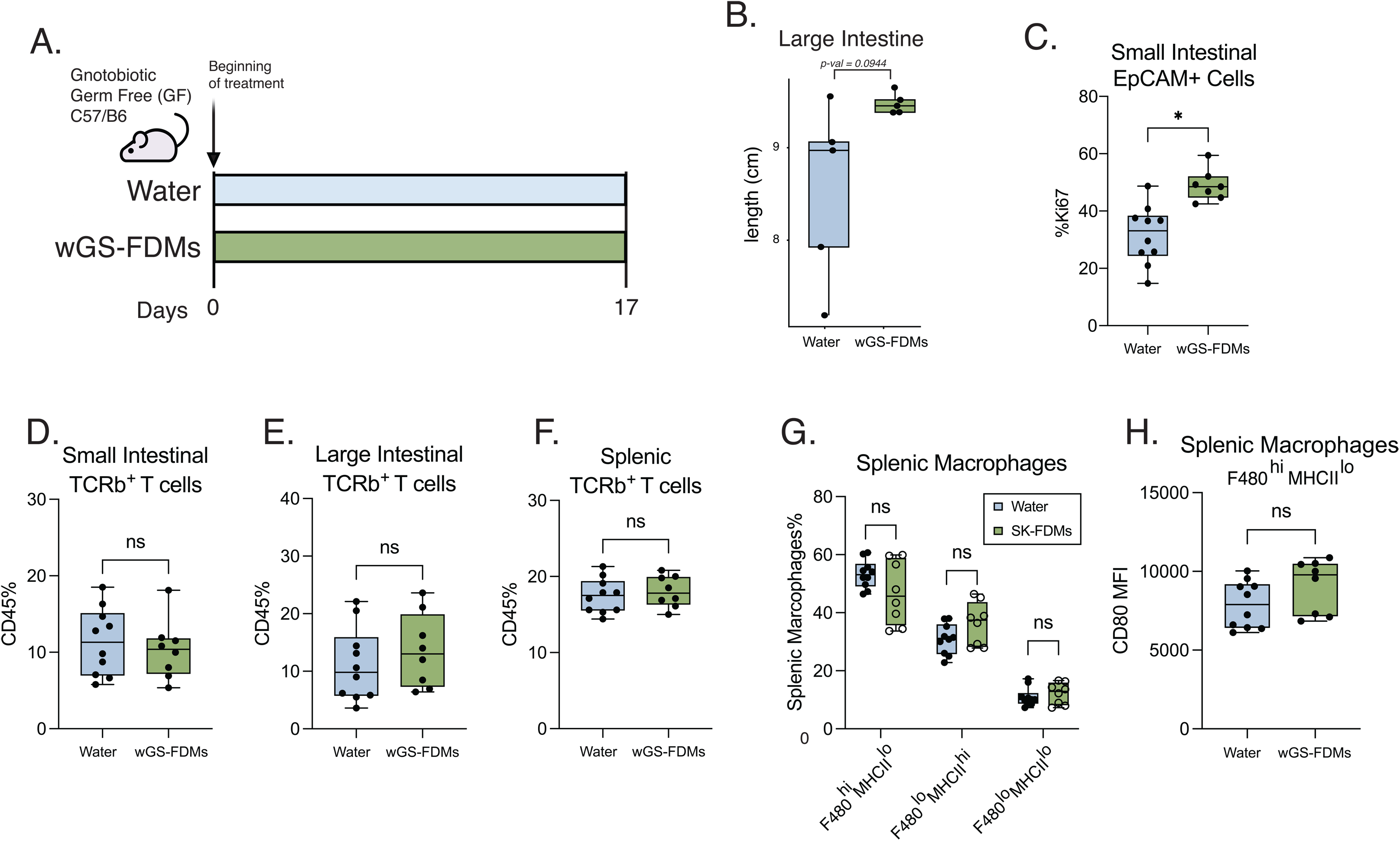
wGS-FDMs promote intestinal epithelial proliferation without altering T cell or macrophage populations in germ-free mice. (A) Schematic of the experimental design. Germ-free (GF) C57BL/6 mice were administered water or wGS-FDMs daily from day 1 through day 17. (B) Large intestine length (cm) of water and wGS-FDM-treated GF mice at endpoint (day 17) (p = 0.0944). (C) Percent of Ki67⁺ cells among EpCAM⁺ small intestinal epithelial cells in water and wGS-FDM-treated mice. (D) Percent of TCRβ⁺ T cells among CD45⁺ cells in the small intestine lamina propria, (E) large intestine lamina propria, and (F) spleen of water and wGS-FDM-treated mice. (G) Proportions of splenic macrophage subsets — F480ʰⁱMHCIIˡᵒ, F480ˡᵒMHCIIʰⁱ, and F480ˡᵒMHCIIˡᵒ — as a percentage of total splenic macrophages in water and wGS-FDM-treated mice. (H) CD80 expression (MFI) on F480ʰⁱMHCIIˡᵒ splenic macrophages from water and wGS-FDM-treated mice. Data are presented as box-and-whisker plots (median ± interquartile range, with whiskers extending to minimum and maximum values) with individual data points overlaid. Statistical comparisons were performed using a two-tailed Mann–Whitney U test or unpaired Student’s t-test, as appropriate. *p < 0.05; ns, not significant. n = 6–10 mice per group.

**Supplemental Figure 10.**
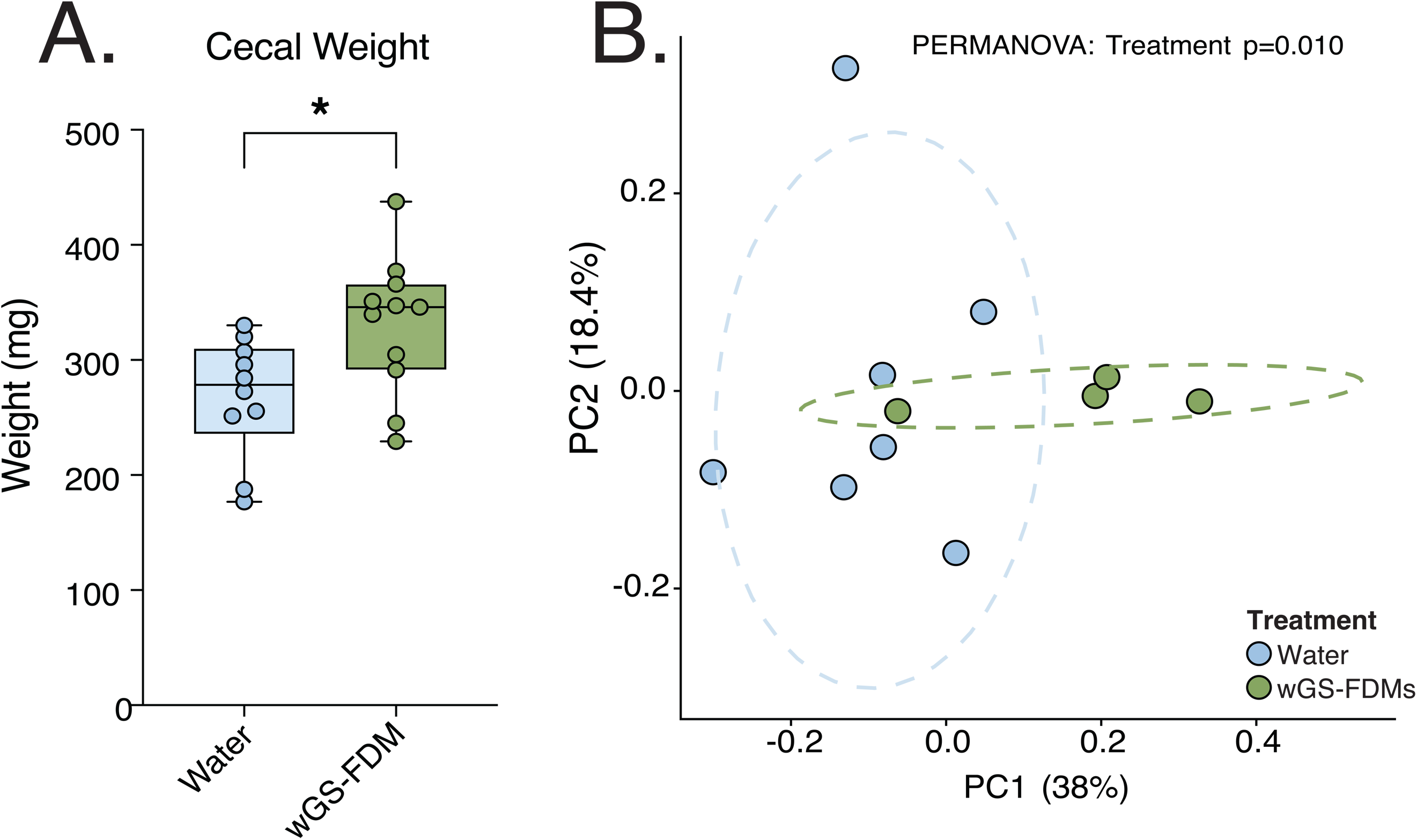
wGS-FDM treatment, in DSS colitis, increases cecal weight and shapes a distinct gut microbial community. (A) Cecal weight (mg) of water and wGS-FDM-treated mice at the final timepoint (day 17). Data are presented as box-and-whisker plots (median ± interquartile range, with whiskers extending to minimum and maximum values) with individual data points overlaid. (B) Principal coordinates analysis (PCoA) of Bray-Curtis dissimilarity of gut microbial communities from water and wGS-FDM-treated mice at the final timepoint (day 17). Each point represents an individual mouse. Dashed ellipses indicate 95% confidence intervals for each treatment group. Treatment groups were significantly separated along PC1 (38% variance explained) and PC2 (18.4% variance explained) (PERMANOVA: F = 2.0021, R² = 0.143, p = 0.010, 999 permutations). Statistical comparison was performed using an unpaired Student’s t-test. *p < 0.05. n = 10–13 mice per group.

**Supplemental Figure 11.**
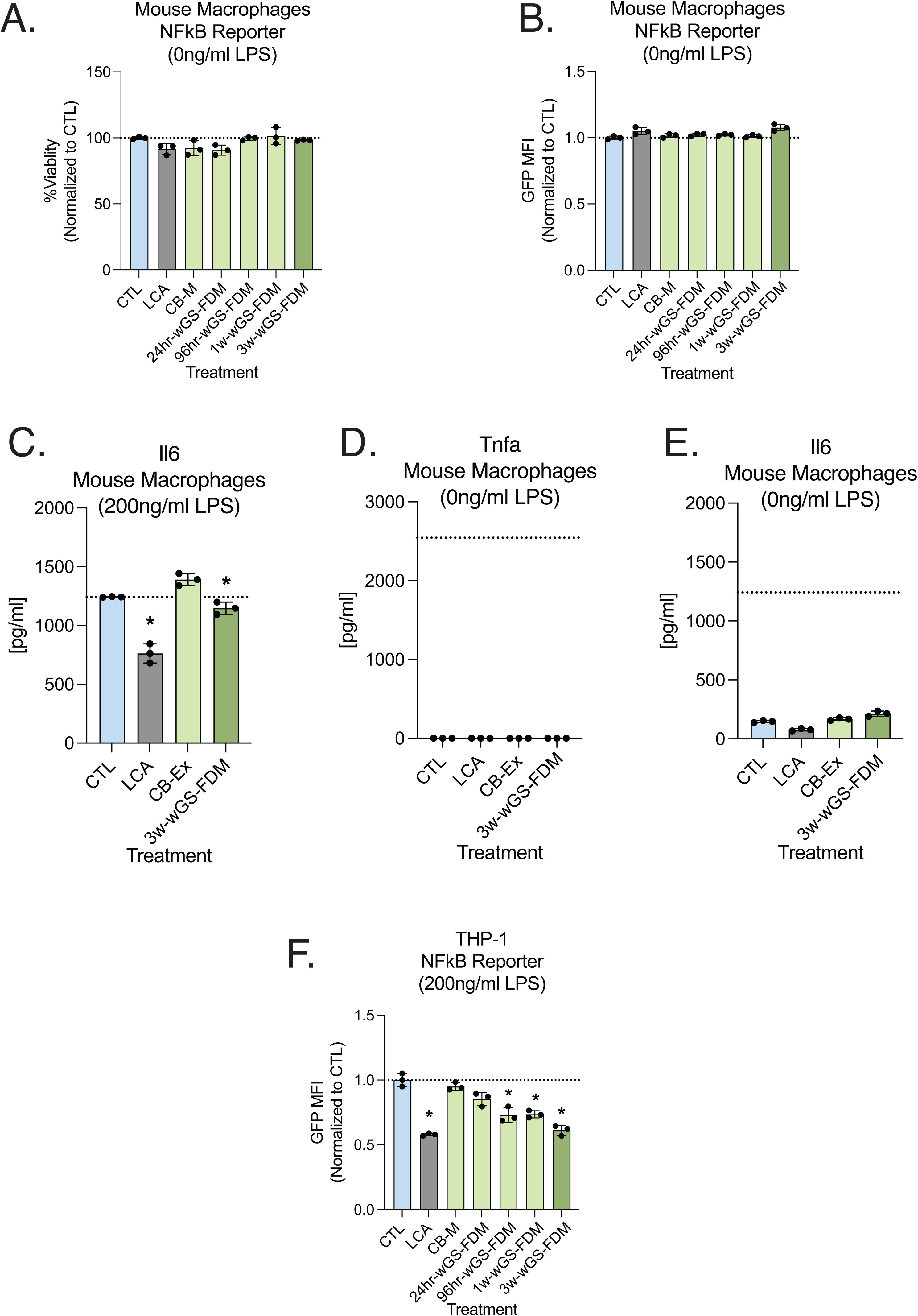
wGS-FDMs suppress NFκB activity and IL-6 secretion in LPS-stimulated macrophages without affecting cell viability or basal inflammatory tone. (A) Percent viability of mouse macrophages expressing an NFκB–GFP reporter, normalized to control (CTL), following treatment with lithocholic acid (LCA), cabbage metabolites (CB-M), 24hr-wGS-FDM, 96hr-wGS-FDM, 1w-wGS-FDM, or 3w-wGS-FDM in the absence of lipopolysaccharide (LPS). (B) NFκB activity, measured as GFP MFI normalized to CTL, in mouse macrophage NFκB reporter cells treated as in (A) under basal conditions (0 ng/ml LPS). (C) IL-6 secretion (pg/ml) by mouse macrophages stimulated with LPS (200 ng/ml) and treated with CTL, LCA, CB-Ex, or 3w-wGS-FDM. (D) TNF-α secretion (pg/ml) by mouse macrophages treated with CTL, LCA, CB-Ex, or 3w-wGS-FDM under basal conditions (0 ng/ml LPS). (E) IL-6 secretion (pg/ml) by mouse macrophages treated with CTL, LCA, CB-Ex, or 3w-wGS-FDM under basal conditions (0 ng/ml LPS). (F) NFκB activity (GFP MFI normalized to CTL) in THP-1 NFκB reporter cells stimulated with LPS (200 ng/ml) and treated with LCA, CB-M, 24hr-wGS-FDM, 96hr-wGS-FDM, 1w-wGS-FDM, or 3w-wGS-FDM. Data are presented as mean ± SD with individual data points overlaid. Dashed lines indicate the CTL reference value. Statistical comparisons were performed using one-way ANOVA with Dunnett’s post hoc test or a Kruskal–Wallis test with Dunn’s correction, as appropriate, comparing each treatment to CTL. *p < 0.05. n = 3–5 independent experiments per group.

**Supplemental Figure 12.**
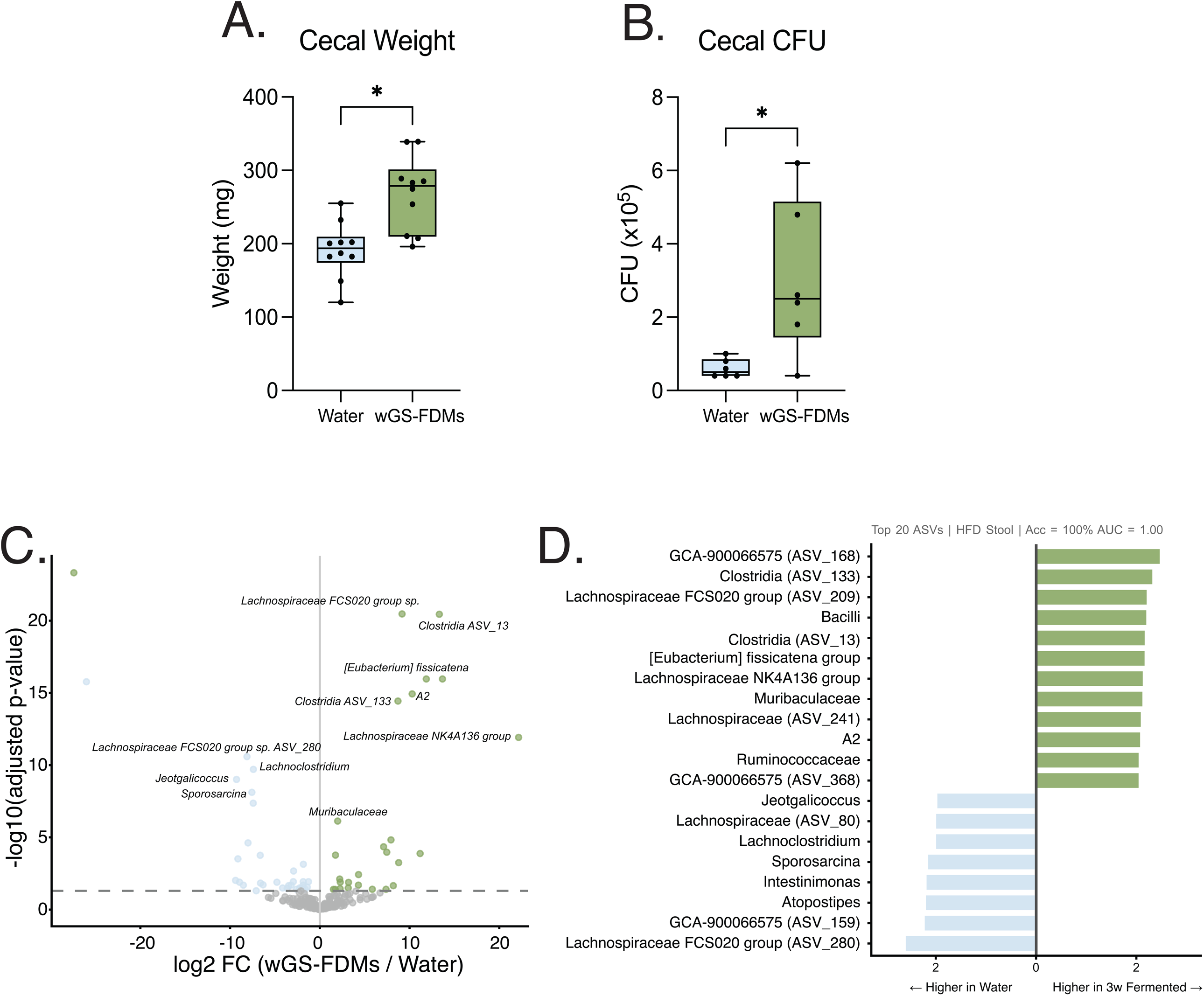
wGS-FDM treatment increases cecal bacterial load and enriches distinct microbial taxa in HFD mice. (A) Cecal weight (mg) of water and wGS-FDM-treated mice at the final timepoint (week 10) presented as box-and-whisker plots (median ± interquartile range, with whiskers extending to minimum and maximum values) with individual data points overlaid. (B) Cecal bacterial burden, quantified as colony-forming units (CFU × 10⁵), from water and wGS-FDM-treated mice presented as box-and-whisker plots as described in (A). (C) Volcano plot depicting differentially abundant amplicon sequence variants (ASVs) between wGS-FDM- and water-treated mice over weeks of treatment. The x-axis shows log2 fold change (wGS-FDMs / Water) and the y-axis shows −log10(adjusted p-value). The dashed horizontal line indicates the significance threshold (adjusted p < 0.05). Green points represent ASVs significantly enriched in wGS-FDM-treated mice; blue points represent ASVs enriched in water-treated mice. Select taxa are labeled. Differential abundance analysis was performed using DESeq2 with Benjamini– Hochberg correction for multiple comparisons. (D) Horizontal bar plot of the top 20 ASVs identified in high-fat diet (HFD) stool, ranked by a classifier with 100% accuracy (AUC = 1.00). Green bars indicate ASVs more abundant in 3-week fermented wGS-FDM-treated mice; blue bars indicate ASVs more abundant in Water-treated mice. Statistical comparisons were performed using an unpaired Student’s t-test. *p < 0.05. n = 8–10 mice per group.

**Supplemental Figure 13.**
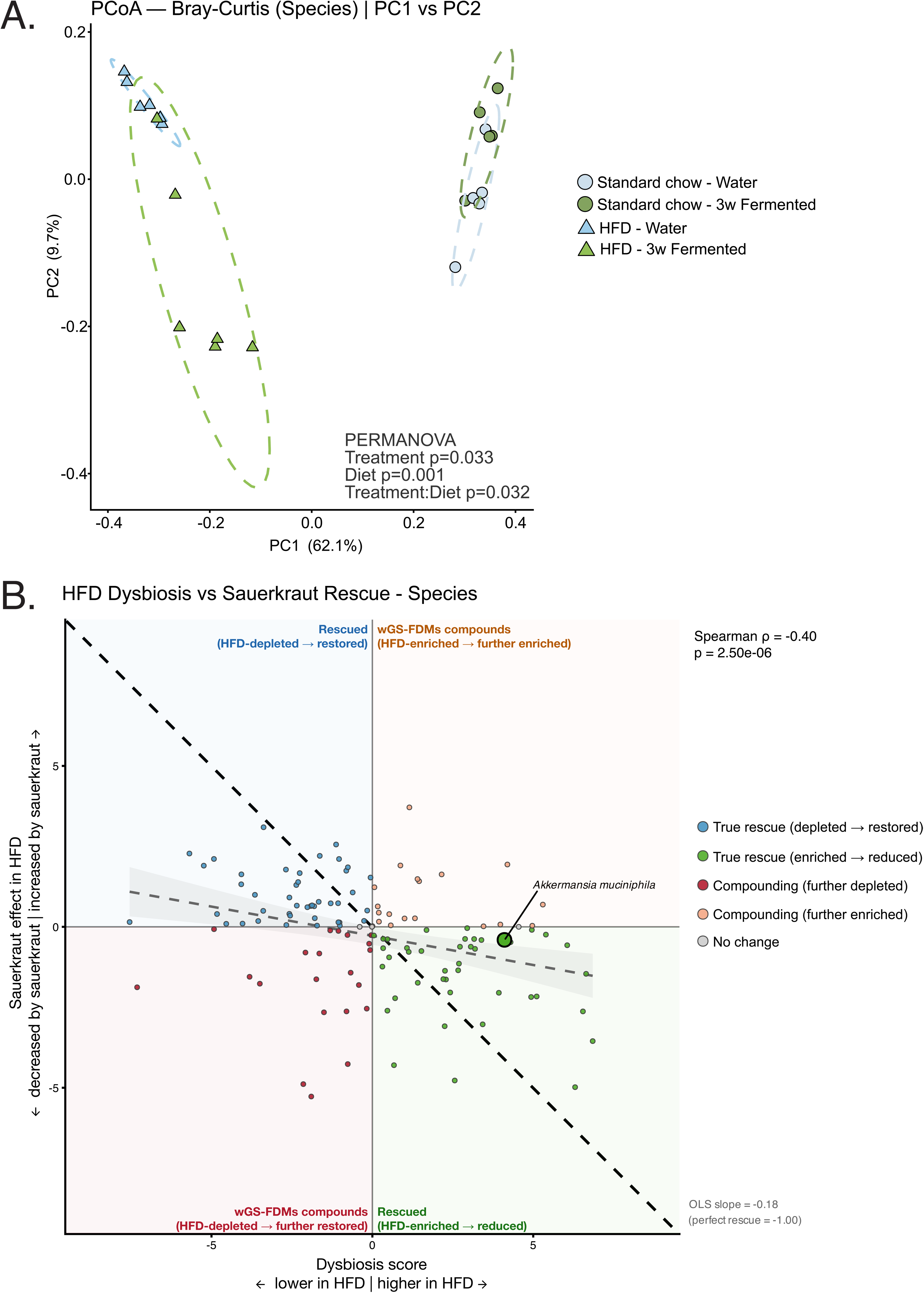
wGS-FDM treatment shifts gut microbiome. (A) Principal coordinates analysis (PCoA) of Bray-Curtis dissimilarity of gut microbial communities from baseline and HFD mice treated with wGS-FDM or untreated at the final timepoint. PERMANOVA revealed significant effects of both diet (p = 0.001) and treatment (p = 0.032). (B) Comparison of HFD-driven dysbiosis score to sauerkraut effect in HFD model. The x-axis indicated the log2 fold-change of HFD water compared to the baseline water group. The y-axis indicated the log2 fold-change of HFD wGS-FDM treatment to HFD water treatment. The black dashed line represents a perfect rescue, while the grey dashed line indicates the observed OLD trend (OLS slope = -0.18). (Spearman ρ = -0.040, p-value = 2.50e-06).

## Citations

1. Sonnenburg, J. L. & Bäckhed, F. Diet-microbiota interactions as moderators of human metabolism. Nature 535, 56–64 (2016).

2. Rooks, M. G. & Garrett, W. S. Gut microbiota, metabolites and host immunity. Nat. Rev. Immunol. 16, 341–352 (2016).

3. Zheng, D., Liwinski, T. & Elinav, E. Interaction between microbiota and immunity in health and disease. Cell Res. 30, 492–506 (2020).

4. Litvak, Y., Byndloss, M. X. & Bäumler, A. J. Colonocyte metabolism shapes the gut microbiota. Science 362, eaat9076 (2018).

5. Salvi, P. S. & Cowles, R. A. Butyrate and the intestinal epithelium: Modulation of proliferation and inflammation in homeostasis and disease. Cells 10, 1775 (2021).

6. Yang, W. & Cong, Y. Gut microbiota-derived metabolites in the regulation of host immune responses and immune-related inflammatory diseases. Cell. Mol. Immunol. 18, 866–877 (2021).

7. Liu, Y. et al. Clostridium sporogenes uses reductive Stickland metabolism in the gut to generate ATP and produce circulating metabolites. Nat. Microbiol. 7, 695–706 (2022).

8. Lopez-Siles, M., Duncan, S. H., Garcia-Gil, L. J. & Martinez-Medina, M. Faecalibacterium prausnitzii: from microbiology to diagnostics and prognostics. ISME J. 11, 841–852 (2017).

9. Dodd, D. et al. A gut bacterial pathway metabolizes aromatic amino acids into nine circulating metabolites. Nature 551, 648–652 (2017).

10. Byndloss, M. X. et al. Microbiota-activated PPAR-γ signaling inhibits dysbiotic Enterobacteriaceae expansion. Science 357, 570–575 (2017).

11. Van Treuren, W. & Dodd, D. Microbial contribution to the human metabolome: Implications for health and disease. Annu. Rev. Pathol. 15, 345–369 (2020).

12. Caffrey, E. B., Sonnenburg, J. L. & Devkota, S. Our extended microbiome: The human-relevant metabolites and biology of fermented foods. Cell Metab. 36, 684–701 (2024).

13. Mukherjee, A., Breselge, S., Dimidi, E., Marco, M. L. & Cotter, P. D. Fermented foods and gastrointestinal health: underlying mechanisms. Nat. Rev. Gastroenterol. Hepatol. 21, 248–266 (2024).

14. Fischbach, M. A. & Sonnenburg, J. L. Eating for two: how metabolism establishes interspecies interactions in the gut. Cell Host Microbe 10, 336–347 (2011).

15. Lavelle, A. & Sokol, H. Gut microbiota-derived metabolites as key actors in inflammatory bowel disease. Nat. Rev. Gastroenterol. Hepatol. 17, 223–237 (2020).

16. David, L. A. et al. Diet rapidly and reproducibly alters the human gut microbiome. Nature 505, 559–563 (2014).

17. Wu, G. D. et al. Linking long-term dietary patterns with gut microbial enterotypes. Science 334, 105–108 (2011).

18. Rothschild, D. et al. Environment dominates over host genetics in shaping human gut microbiota. Nature 555, 210–215 (2018).

19. Sonnenburg, J. L. & Sonnenburg, E. D. Vulnerability of the industrialized microbiota. Science 366, eaaw9255 (2019).

20. Carter, M. M. et al. Ultra-deep sequencing of Hadza hunter-gatherers recovers vanishing gut microbes. Cell 186, 3111–3124.e13 (2023).

21. Zhernakova, A. et al. Population-based metagenomics analysis reveals markers for gut microbiome composition and diversity. Science 352, 565–569 (2016).

22. Manor, O. et al. Health and disease markers correlate with gut microbiome composition across thousands of people. Nat. Commun. 11, 5206 (2020).

23. Vangay, P. et al. US Immigration Westernizes the Human Gut Microbiome. Cell 175, 962–972.e10 (2018).

24. Yatsunenko, T. et al. Human gut microbiome viewed across age and geography. Nature 486, 222–227 (2012).

25. Smillie, C. S. et al. Strain Tracking Reveals the Determinants of Bacterial Engraftment in the Human Gut Following Fecal Microbiota Transplantation. Cell Host Microbe 23, 229–240.e5 (2018).

26. Wilson, B. C., Vatanen, T., Cutfield, W. S. & O’Sullivan, J. M. The super-donor phenomenon in fecal Microbiota transplantation. Front. Cell. Infect. Microbiol. 9, 2 (2019).

27. Suez, J. et al. Post-antibiotic gut mucosal microbiome reconstitution is impaired by probiotics and improved by autologous FMT. Cell 174, 1406–1423.e16 (2018).

28. Kristensen, N. B. et al. Alterations in fecal microbiota composition by probiotic supplementation in healthy adults: a systematic review of randomized controlled trials. Genome Med. 8, 52 (2016).

29. Wastyk, H. C. et al. Gut-microbiota-targeted diets modulate human immune status. Cell 184, 4137–4153.e14 (2021).

30. Jakubczyk, K., Kałduńska, J., Kochman, J. & Janda, K. Chemical profile and antioxidant activity of the kombucha beverage derived from white, green, black and red tea. Antioxidants (Basel) 9, 447 (2020).

31. Kasperek, M. C. et al. Microbial aromatic amino acid metabolism is modifiable in fermented food matrices to promote bioactivity. bioRxiv (2023) doi:10.1101/2023.12.21.572869.

32. Wei, L. & Marco, M. L. The fermented cabbage metabolome and its protection against cytokine-induced intestinal barrier disruption of Caco-2 monolayers. Appl. Environ. Microbiol. 91, e0223424 (2025).

33. Li, Y. et al. Fermentation of celery (Apium graveolens L.) with Lactobacillus plantarum NCU116: Impact on physicochemical properties, free amino acids, and volatile aroma compounds. Food Biosci. 68, 106680 (2025).

34. Chen, W., Zhang, L. & Li, S. Metagenomics and untargeted metabolomics analyses to unravel the formation mechanism of characteristic metabolites in mung bean sour liquid during different fermentation stages. Lebenson. Wiss. Technol. 233, 118488 (2025).

35. Nielsen, E. S. et al. Lacto-fermented sauerkraut improves symptoms in IBS patients independent of product pasteurisation - a pilot study. Food Funct. 9, 5323–5335 (2018).

36. Schropp, N. et al. The impact of regular sauerkraut consumption on the human gut microbiota: a crossover intervention trial. Microbiome 13, 52 (2025).

37. Miller, E. R., O’Mara Schwartz, J., Cox, G. & Wolfe, B. E. A gnotobiotic system for studying microbiome assembly in the phyllosphere and in vegetable fermentation. J. Vis. Exp. (2020) doi:10.3791/61149.

38. Miller, E. R. et al. Establishment limitation constrains the abundance of lactic acid bacteria in the Napa cabbage phyllosphere. Appl. Environ. Microbiol. 85, (2019).

39. Han, S. et al. A metabolomics pipeline for the mechanistic interrogation of the gut microbiome. Nature 595, 415–420 (2021).

40. Han, S., Guiberson, E. R., Li, Y. & Sonnenburg, J. L. High-throughput identification of gut microbiome-dependent metabolites. Nat. Protoc. 19, 2180–2205 (2024).

41. Roager, H. M. & Licht, T. R. Microbial tryptophan catabolites in health and disease. Nat. Commun. 9, 3294 (2018).

42. Agus, A., Planchais, J. & Sokol, H. Gut Microbiota regulation of tryptophan metabolism in health and disease. Cell Host Microbe 23, 716–724 (2018).

43. Shin, H.-K. & Bang, Y.-J. Aromatic amino acid metabolites: Molecular messengers bridging immune-Microbiota communication. Immune Netw. 25, e10 (2025).

44. Liu, Y., Hou, Y., Wang, G., Zheng, X. & Hao, H. Gut microbial metabolites of aromatic amino acids as signals in host-microbe interplay. Trends Endocrinol. Metab. 31, 818– 834 (2020).

45. Kilstrup, M., Hammer, K., Ruhdal Jensen, P. & Martinussen, J. Nucleotide metabolism and its control in lactic acid bacteria. FEMS Microbiol. Rev. 29, 555–590 (2005).

46. Li, V. L. et al. An exercise-inducible metabolite that suppresses feeding and obesity. Nature 606, 785–790 (2022).

47. Li, Z. et al. Dietary butyrate ameliorates metabolic health associated with selective proliferation of gut Lachnospiraceae bacterium 28-4. JCI Insight 8, e166655 (2023).

48. Functional and Genomic Variation between HumanDerived Isolates of Lachnospiraceae Reveals Interand Intra-Species Diversity.

49. Riva, A. et al. Identification of inulin-responsive bacteria in the gut microbiota via multi-modal activity-based sorting. Nat. Commun. 14, 8210 (2023).

50. Windeløv, J. A. et al. Why is it so difficult to measure glucagon-like peptide-1 in a mouse? Diabetologia 60, 2066–2075 (2017).

51. Smits, M. M. et al. In vivo inhibition of dipeptidyl peptidase 4 allows measurement of GLP-1 secretion in mice. Diabetes 73, 671–681 (2024).

52. Gagnon, J. & Brubaker, P. L. NCI-H716 Cells. in The Impact of Food Bioactives on Health 221–228 (Springer International Publishing, Cham, 2015).

53. Spencer, S. P., Fragiadakis, G. K. & Sonnenburg, J. L. Pursuing human-relevant gut Microbiota-immune interactions. Immunity 51, 225–239 (2019).

54. Matsumoto, M., Kurihara, S., Kibe, R., Ashida, H. & Benno, Y. Longevity in mice is promoted by probiotic-induced suppression of colonic senescence dependent on upregulation of gut bacterial polyamine production. PLoS One 6, e23652 (2011).

55. Smith, B. J. et al. Changes in the gut microbiome and fermentation products concurrent with enhanced longevity in acarbose-treated mice. BMC Microbiol. 19, 130 (2019).

56. Derrien, M., Belzer, C. & de Vos, W. M. Akkermansia muciniphila and its role in regulating host functions. Microb. Pathog. 106, 171–181 (2017).

57. Ghotaslou, R. et al. The metabolic, protective, and immune functions of Akkermansia muciniphila. Microbiol. Res. 266, 127245 (2023).

58. Rescigno, M. The intestinal epithelial barrier in the control of homeostasis and immunity. Trends Immunol. 32, 256–264 (2011).

59. van Wijk, F. & Cheroutre, H. Intestinal T cells: facing the mucosal immune dilemma with synergy and diversity. Semin. Immunol. 21, 130–138 (2009).

60. Nagler-Anderson, C. Man the barrier! Strategic defences in the intestinal mucosa. Nat. Rev. Immunol. 1, 59–67 (2001).

61. Inaba, K. et al. The formation of immunogenic major histocompatibility complex class II-peptide ligands in lysosomal compartments of dendritic cells is regulated by inflammatory stimuli. J. Exp. Med. 191, 927–936 (2000).

62. Linsley, P. S. & Ledbetter, J. A. The role of the CD28 receptor during T cell responses to antigen. Annu. Rev. Immunol. 11, 191–212 (1993).

63. Kurotaki, D., Uede, T. & Tamura, T. Functions and development of red pulp macrophages: Biology of red pulp macrophages. Microbiol. Immunol. 59, 55–62 (2015).

64. Postler, T. S. & Ghosh, S. Understanding the holobiont: How microbial metabolites affect human health and shape the immune system. Cell Metab. 26, 110–130 (2017).

65. Smith, P. M. et al. The microbial metabolites, short-chain fatty acids, regulate colonic Treg cell homeostasis. Science 341, 569–573 (2013).

66. Furusawa, Y. et al. Commensal microbe-derived butyrate induces the differentiation of colonic regulatory T cells. Nature 504, 446–450 (2013).

67. Koh, A., De Vadder, F., Kovatcheva-Datchary, P. & Bäckhed, F. From dietary fiber to host physiology: Short-chain fatty acids as key bacterial metabolites. Cell 165, 1332– 1345 (2016).

68. Yu, A. I. et al. Gut Microbiota modulate CD8 T cell responses to influence colitis-associated tumorigenesis. Cell Rep. 31, 107471 (2020).

69. Vacca, M. et al. The controversial role of human gut Lachnospiraceae. Microorganisms 8, 573 (2020).

70. Cani, P. D., Everard, A. & Duparc, T. Gut microbiota, enteroendocrine functions and metabolism. Curr. Opin. Pharmacol. 13, 935–940 (2013).

71. Zeng, Y., Wu, Y., Zhang, Q. & Xiao, X. Crosstalk between glucagon-like peptide 1 and gut microbiota in metabolic diseases. MBio 15, e0203223 (2024).

72. Thomas, C. et al. TGR5-mediated bile acid sensing controls glucose homeostasis. Cell Metab. 10, 167–177 (2009).

73. Chao, J., Coleman, R. A., Keating, D. J. & Martin, A. M. Gut microbiome regulation of gut hormone secretion. Endocrinology 166, (2025).

74. Daniel, N. et al. Gut microbiota and fermentation-derived branched chain hydroxy acids mediate health benefits of yogurt consumption in obese mice. Nat. Commun. 13, 1343 (2022).

## Methods References

1. Han, S., Guiberson, E. R., Li, Y. & Sonnenburg, J. L. High-throughput identification of gut microbiome-dependent metabolites. Nat. Protoc. 19, 2180–2205 (2024).

2. Sinha, S. R. et al. Dysbiosis-induced secondary bile acid deficiency promotes intestinal inflammation. Cell Host Microbe 27, 659–670.e5 (2020).

3. Sonnenburg, E. D. et al. Diet-induced extinctions in the gut microbiota compound over generations. Nature 529, 212–215 (2016).

4. Goodyear, A. W., Kumar, A., Dow, S. & Ryan, E. P. Optimization of murine small intestine leukocyte isolation for global immune phenotype analysis. J. Immunol. Methods 405, 97–108 (2014).

5. Covarrubias, S. et al. CRISPR/Cas-based screening of long non-coding RNAs (lncRNAs) in macrophages with an NF-κB reporter. J. Biol. Chem. 292, 20911–20920 (2017).

6. Halasz, H. et al. CRISPRi screens identify the lncRNA, LOUP, as a multifunctional locus regulating macrophage differentiation and inflammatory signaling. Proc. Natl. Acad. Sci. U. S. A. 121, e2322524121 (2024).

