## Supplementary Table 5 for "Fermentation-Derived Metabolites Shape Host Biology to Attenuate Severity of Inflammatory and Metabolic Disease"

Supplemental Table 5: Antibodies used in flow cytometry experiments

| Marker | Manufacture | Catalog ID | Fluorophore | Clone |
|---|---|---|---|---|
| Live Dead | Invitrogen | L23105 | LIVE/DEAD Blue | |
| CD45 | Biolegend | 103130 | PerCP | 30-F11 |
| B220 | BD Horizon™ | BDB612972 | BUV661 | RA3-6B2 |
| NK1.1 (CD161) | Biolegend | 108724 | APC-Cy7 | PK136 |
| TCRb | Biolegend | 109224 | AF700 | H57-597 |
| CD90.2 | BD Horizon™ | 741909 | BUV805 | 30-H12 |
| CD8a | Biolegend | 100779 | SparkBlue 550 | 53-6.7 |
| CD4 | Biolegend | 100476 | Spark NIR 685 | GK1.5 |
| Foxp3 | eBioscience | 48-5773-82 | ef450 | FJK-16s |
| Rorgt | eBioscience | 12-6981-82 | PE | B2D |
| Tbet | eBioscience | 25-5825-82 | PE-Cy7 | eBio4B10(4B10) |
| F480 | Biolegend | 123108 | FITC | BM8 |
| Ki67 | BD Biosciences | BDB563755 | BV711 | B56 |
| EpCam (CD326) | Biolegend | 118217 | APC | G8.8 |
| Siglec-F (CD170) | Biolegend | 155525 | PerCP Cy5.5 | S17007L |
| Ly6G | BD Biosciences | 612921 | BUV563 | 1A8 |
| Ly6C | Biolegend | 128041 | BV785 | HK1.4 |
| CD11b | Biolegend | 101233 | BV570 | M1/70 |
| CD11c | Biolegend | 117312 | AF647 | N418 |
| I-A/I-E | Biolegend | 107620 | Pacific Blue | M5/114.15.2 |
| CD103 | Biolegend | 121421 | BV421 | 2E7 |
| TCRgd | eBioscience | 15-5711-81 | PE-Cy5 | eBioGL3 (GL-3, GL3) |
| CD44 | Biolegend | 103082 | PerCP-Fire806 | IM7 |
| CD62L | BD Horizon™ | 563117 | BV510 | MEL-14 |
| CD86 | BD Biosciences | BDB750437 | BUV496 | PO2 (RUO) |
| CD206 | Biolegend | 141721 | BV605 | C068C2 |
| CD80 | Biolegend | 104738 | PE Dazzle 594 | 16-10A1 |
